# Weak Vestibular Response in Persistent Developmental Stuttering: Implications for Own Voice Identification

**DOI:** 10.1101/2020.11.24.396283

**Authors:** Max Gattie, Elena Lieven, Karolina Kluk

## Abstract

Speech-motor and psycholinguistic models employ feedback control from an auditory stream corresponding to own voice. Such models underspecify how own voice is identified. It is proposed that own voice is identified through coincidence detection between the neural firing rates arising from deflection of cochlear and vestibular mechanoreceptors by the sound and vibration generated during vocalisation. The coincidence detection is proposed to differ in people who stutter. In an update to the approach-avoidance conflict model of Sheehan (1953, 1975) instances of stuttering are proposed to coincide with uncertainty over an ongoing speech act. Discussion covers speech-induced suppression, auditory scene analysis, and theories of mental content.

## 1. Introduction

Speech-motor and psycholinguistic models describe a feedforward system in which articulatory muscles receive coordinated nerve impulses with sufficient detail to generate speech sounds (e.g. Hickok & Poeppel, 2007; Levelt et al., 1999; Tourville & Guenther, 2011). Typically they employ feedback control as a check for error (Helmholtz, 1886; von Holst & Mittelstädt, 1950; Fairbanks, 1954). Predictive feedback control avoids instability due to timing delay by checking for sensory error against a forward model of the speech-motor plan (see review in Parrell & Houde, 2019). Errors checked for might include articulatory malfunction, or mismatch between spoken and intended message – the nature of the error checked for will vary, depending on the nature of the model.

Such models underspecify how an auditory stream corresponding to own voice is identified (i.e. an auditory stream defined as per Bregman, 1990). A typical requirement is that a mental representation of expected auditory consequences is referred to, or is already identical with, an auditory target map (O’Callaghan, 2015). The question arises of how such reference is managed in the opposite direction – how an auditory target map for own voice is created from ambient sound and vibration.

Greater understanding of own voice identification could improve speech-motor and psycholinguistic models. For example, previously overlooked activity in the auditory brainstem and periphery may explain otherwise intractable difficulties in understanding the cerebral and cerebellar activity accompanying speech and language. Such an approach is taken in the current article. A hypothesis is formulated for own voice identification. The hypothesis is then developed to provide an account of stuttering, a DSM-V diagnosis characterised by involuntary prolongations and repetitions during speech.

The article will proceed as follows. Section 2 will describe the hypothesis of own voice identification. Section 3 will build on the hypothesis of Section 2 to present a novel account of stuttering, REMATCH (Reflexivity and Communicative Mismatch). Section 4 will provide discussion of themes arising from sections 2 and 3. In this way, the article will extend from a biophysical account of own voice identification, to a psychosocial account of interpersonal communication. It will progress from audiology, to speech-motor theory, to psycholinguistics and social psychology.

Hypothesis formulation follows inference to the best explanation (Lipton, 2004). Best explanation arguments are mutually supportive. In other words, if one has a best explanation argument of T, and one has a best explanation argument of D, it follows that one has a best explanation argument of (T + D). This pertains even if D is partially reliant on T. This system (sometimes referred to as abduction) differs from, for example, multiplicative combination of probabilities in which the combined probability is lower than either of its constituents. Refuting a best explanation argument requires presentation of a better explanation. The discussion in Section 4 will summarise the scope of the best explanation argument. To aid that discussion, hypotheses will be presented following the Methodology of Scientific Research Programmes described by Lakatos (1970). This refers to a “hard core” of (generally unfalsifiable) hypotheses, along with a “protective belt” of testable auxiliary hypotheses. Distinction will also be made between the two kinds of causal explanation described by Botterill (2010). Process explanations are of how something happens, whereas contrastive explanations are of why something happens. These two kinds of explanation interact as understanding of causation is acquired and enhanced.

## 2. Hypothesis of Own Voice Identification

### 2.1 Explanatory target

Own voice identification is a specific instance of the cocktail party problem (Bee & Micheyl, 2008), an outstanding issue in auditory scene analysis in which there is no principled basis for discrimination in a multi-talker scenario. It is an example of an ill-posed problem (Hadamard 1902, 1923; Poggio & Koch, 1985), sometimes referred to as an inverse problem, in which there is no mathematically unique solution.

### 2.2 Candidate explanations

There is no prior research offering a basis by which an own voice auditory stream is specifically distinguished from ambient sound and vibration (Shamma & Micheyl, 2010; Remez & Thomas, 2013; Bronkhorst, 2015). The most closely related literature emphasises the importance of body conducted vibration during own speech (von Békésy, 1949; Maurer & Landis, 1990; Pörschmann, 2000; Sohmer & Freeman, 2001; Shuster & Durrant, 2003; Reinfeldt et al., 2010; Meekings et al., 2015) or else describes self talk and private speech through a Vygotskian developmental perspective (e.g. Fernyhough & Russell, 1997; Atencio & Montero, 2009; Lupyan & Swingley, 2012).

There is also a large body of work about the role of own voice in speech monitoring systems (e.g. Postma, 2000; Buschbaum, 2001; Ozdemir et al., 2007; Huettig & Hartsuiker, 2010; Nozari et al., 2011; Lind et al., 2014; Acheson & Hagoort, 2014; Kröger et al., 2016) or sensory-motor integration (e.g. Jürgens, 2002; Kaplan et al., 2008; Rosa et al., 2008; Zheng et al., 2010; Hickok et al., 2011; Behroozmand et al., 2015; Houde et al., 2015). This literature takes as a starting point that own voice has already been identified as an ascending auditory stream. It therefore does not address the current explanatory target. Literature concerning sensory-motor integration, and in particular the hypothesis of speech-induced suppression, will be discussed in section 2.4.1.

### 2.3 A Novel Hypothesis of Own Voice Identification

#### 2.3.1 Introduction

The nature of the speech auditory brainstem response (BinKhamis et al., 2019) suggests that neural activity corresponding to identification of own voice could occur in the auditory brainstem. The auditory brainstem is innervated through the VIII cranial nerve, from bipolar ganglion cells which interface with mechanoreceptors of the inner ear. Neural activity corresponding to own voice could occur at the earliest within the bipolar ganglion cells of the ear itself.

Inner ear structure is common across mammals, consisting of an osseous labyrinth lined with sensory epithelium, and with several chambers. One of the chambers is the cochlea, a coiled structure containing mechanoreceptors which are deflected by ambient sound frequencies ranging from 20 Hz – 20,000 Hz in humans (Manley & Gummer, 2017). Other chambers comprise the vestibular system. These chambers include semicircular canals, in which mechanoreceptors are deflected by changes in angular velocity. There are also gravitoinertial otoliths, arranged such that mechanoreceptors are deflected by changes in linear velocity, and with resting state deflection corresponding to head orientation (Goldberg, 2012).

The traditional discrimination just described, of cochlear and vestibular chambers into hearing and equilibrial functions, is misleading (Tait, 1932). As for other vertebrates, mammalian otolithic receptors are deflected by vibration as well as by changes in body velocity or orientation relative to a fixed gravitational field. The vestibular system in mammals responds to vibrational frequencies up to 1,000 Hz, and may phase lock to higher frequencies (Curthoys et al., 2019).

Vestibular sensitivity is considerably greater to vibrations conducted through the body (BC) than to sound waves in air (AC). Electrophysiological studies show that when human responses of vestibular origin are referenced to a 60 dBA sound level typical of conversational speech, AC thresholds are 10 dB above baseline and BC thresholds 25 dB below baseline (McNerney & Burkard, 2011; Welgampola, Rosengren, Halmagyi & Colebatch, 2003). The act of speaking will deflect vestibular mechanoreceptors in humans (Todd, Rosegren & Colebatch, 2008; Curthoys, 2017; Curthoys et al., 2019).

#### 2.3.2 Concurrency Hypothesis

The core hypothesis is that own voice is identified as an auditory stream through coincidence detection between vestibular and cochlear afferents. This will henceforth be referred to as the Concurrency Hypothesis.

The Concurrency Hypothesis describes a biologically grounded mechanism. The biological grounding is that there are two sets of mechanoreceptors for own voice. Figure 1 gives an overview of relevant details. Sound and vibrational energy deflecting sterocilia in cochlear hair cells corresponds to own voice mixed with ambient environmental sounds. Concurrently, vibrational energy deflecting stereocilia in vestibular hair cells corresponds to own voice in isolation. Comparison of nerve impulses arising from cochlear and vestibular mechanoreceptors therefore provides a principled distinction between self and environment.

**Figure 1:**
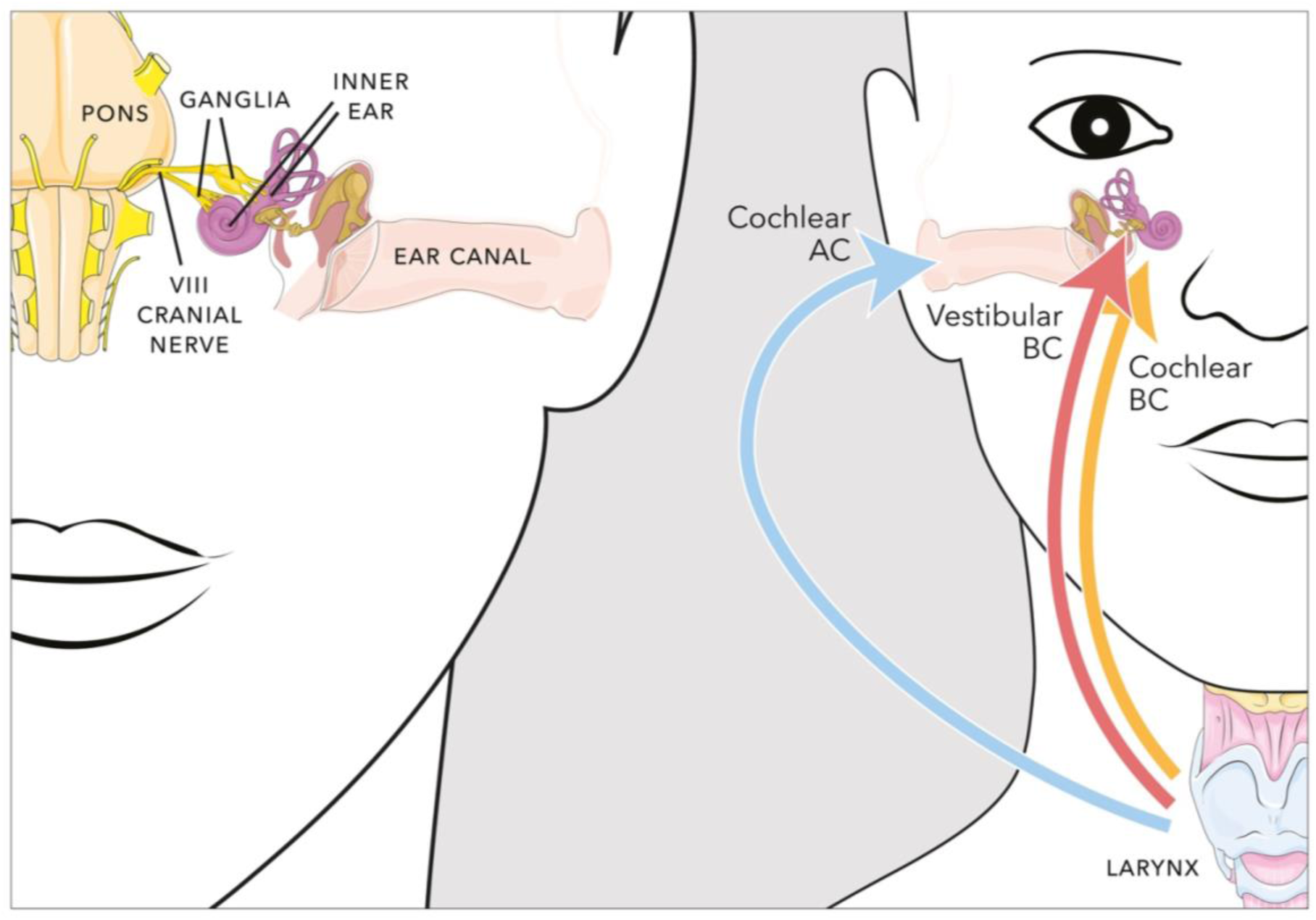
Parameters affecting sound and vibration detection in the human ear. In vivo measurements are difficult, and estimates here are derived from primary sources where possible. For general background on sound source perception, see Yost et al. (2008); for hair cells see Eatock et al. (2006); for voice production see Titze (1994); and for propagation of sound and vibration see Fahy & Thompson (2015). Left hand side: Anatomical parameters. Dendrons of bipolar ganglion cells terminate on sensory epithelial hair cells in the inner ear. Axons from the ganglia project or branch through the VIII cranial nerve to nuclei of the pons and medulla, and (for some axons from vestibular ganglia) the cerebellum. Sensory hair cells fire continuously, with changes in firing rate following deflections due to sound, vibration and movement. Changes in firing rate will in turn modify long-term potentiation of brainstem and cerebellar nerve cells. Right hand side: Acoustic and vibrational parameters. During vocalisation, sound and vibration energy originates predominantly at the larynx (and occasionally higher in the vocal tract; Titze, 1994). Energy propagates via two routes to each ear: air conduction (AC) through air surrounding the head, or body conduction (BC) through the neck and head. The inner ear includes cochlear and vestibular sensory hair cells. Sounds are perceived when AC and BC stimulation above hearing threshold (by definition zero dB HL or higher) deflects stereocilia in cochlear hair cells, opening mechanically gated ion channels which set off a chain of activity culminating in release of neurotransmitters, which in turn will raise potentials in dendrites of ganglion cells belonging to the VIII cranial nerve. Deflection of stereocilia in vestibular hair cells requires a considerably higher stimulus level than that for sterocilia in cochlear hair cells. Welgampola et al. (2003) established electrophysiological vestibular thresholds (VEMPs) at sound levels, as defined at the cochlea, of 31 dB HL for BC stimulation, and 87 dB HL for AC stimulation. Even after adjusting for temporal integration with the brief duration stimuli used in electrophysiological testing, AC vestibular thresholds are 10 dB above, and BC vestibular thresholds 25 dB below, the 60 dBA sound level typical of conversational speech (McNerney & Burkard, 2011). Thus, own voice is either not detected or is very weakly detected via an AC vestibular route. Whereas, unless using alaryngeal speech such as whispering, own voice will consistently be detected by a BC vestibular route. This BC vestibular audition of own voice will persist even if AC and BC cochlear audition of own voice is masked. © Adapted by Max Gattie from illustrations by Servier Medical Art, https://smart.servier.com. Creative Commons 3.0 licence.

Estimating arrival times for own voice stimuli at the inner ear requires consideration of propagation routes (figure 2). Air-conducted (AC) sound can be direct (dAC) or reflected (rAC), whereas body-conducted (BC) vibration can be considered as direct only. Table 1 estimates arrival time at the inner ear at approximately 0.5 ms after vocalisation for both dAC sound and BC vibration. At 60 dBA stimulus levels (typical of vocalisation) BC vibration deflects both cochlear and vestibular mechanoreceptors (McNerney & Burkard, 2011; Welgampola, Rosengren, Halmagyi & Colebatch, 2003). Table 1 compares the propagation timings. Binaural coincidence detection across cochlear and vestibular mechanoreceptors, based on dAC sound and BC vibration, would identify own voice.

**Figure 2:**
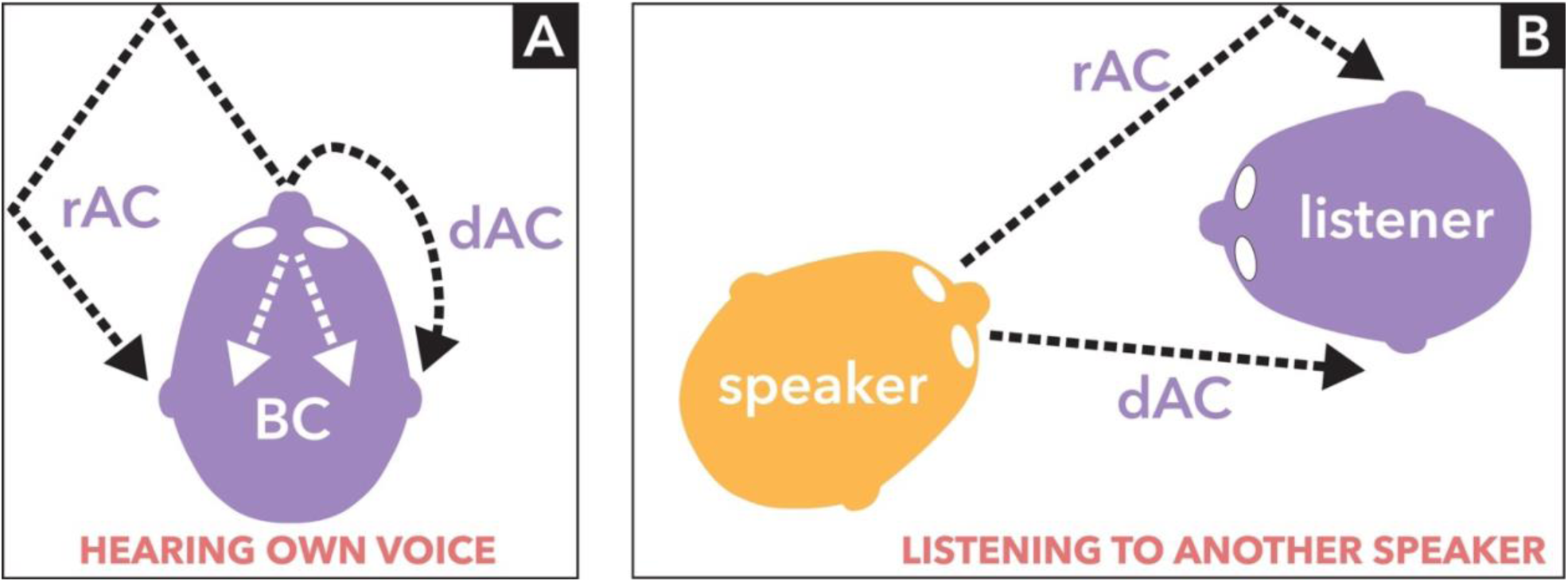
Sound and vibration routes to the ear. Propagation routes are difficult to measure in vivo, and estimates here are derived from primary sources where possible. For general background on sound source perception, see Yost et al. (2008); and for propagation of sound and vibration see Fahy & Thompson (2015). Air-conducted sound is split between reflected (rAC) and direct (dAC) routes (Cabrera et al., 2009; Traer & McDermott, 2016). These are shown in a simplified version. The rAC consists of many environmental reflections with comb filtering (frequencies attenuated or reinforced due to phase differences) as sound energy reaches the ear (Yadav et al., 2012; Arend et al., 2017). The many possible routes for rAC reflect the relationship between body and environment. If reflections of reflections are present (e.g. standing waves inside a room) rAC becomes reverberation. The dAC route is transmitted directly through the air around the speaker’s head. This route includes body reflection, such as that from the shoulders. There is just one form of dAC, which will tend to be stable over the short-term (unless it is windy) and medium-term (unless the head rotates relative to the torso). Conditions in which dAC is unstable tend to also be ones in which conversation is difficult. (A) Transmission time estimates are based on human head dimensions, and will vary according to skull size and individual physiology. When hearing own voice, dAC sound is transmitted at 340 m/s and so will reach the ear in about 0.5 ms. Body conduction (BC) is through bone or soft tissue (Sohmer, 2017; Chordekar et al., 2018). Propagation routes are complex and frequency dependent, will differ between individuals, and have a nature not fully determined in vivo. However, the complexity of propagation routes will be stable in adults, changing only gradually with head composition and body profile across the lifespan. A propagation rate of 300 m/s is likely in humans (Hotehama & Nakagawa, 2012). If so, BC transmission time can be estimated as similar to the 0.5 ms for dAC. A distance of 1.5 cm between cochlear and vestibular hair cells (Ekdale, 2013) gives propagation time for vibration across the inner ear as 0.05 ms. This becomes an upper limit for arrival time difference from a laryngeal source, meaning BC arrival time is coincident to less than 0.05 ms for vestibular and cochlear mechanoreceptors. Routes to the ear for rAC will typically take 2–20 ms (depending on environmental parameters), and will be considerably less stable than for dAC or BC given that the environment, and the position of the head relative to surroundings, can be expected to change continuously. (B) When listening to another speaker, dAC sound energy travelling a direct route between interlocutors is heard first. Energy travelling the longer, indirect route of rAC trails dAC slightly (e.g. by 5–10 ms, depending on environment). Thus, changes in firing rates of inner ear hair cells due to a typical 200 ms CV speech syllable travelling dAC and rAC routes will be spread over a further 2–50 ms or more, depending on proximity of interlocuters and environmental reflections. This overlaps with the time window for the Haas, or precedence, effect – a psychoacoustic phenomenon in which sounds separated by less than about 50 ms are perceptually integrated, with longer delays perceived as echo (Haas, 1951; Wallach et al., 1949). Overwhelmingly, dAC and rAC will have different presentations at each ear, along with comb filtering interactions, such that source localisation is via stereo combination following the duplex theory of Rayleigh (1907). There is in principle a confound for sound sources occupying the “cone of confusion” (a set of points equidistant from each ear) in symmetrical environments or those, like an anechoic chamber, with minimal rAC. In practice such a situation is so unlikely to be sustained that it would not normally have developmental impact (but see Cody et al., 1996). For animals with a pinna, filtering effects of the pinna reduce localisation inaccuracy for sources within the cone of confusion (Musican & Butler, 1984). © Creative Commons 4.0 licence.

**Table 1:** Sound and vibrational energy is transmitted to each ear through body conduction (BC) and direct and reflected air conduction (dAC/rAC), and can deflect two sets of mechanoreceptors in each inner ear. At stimulus levels typical of own voice, vestibular mechanoreceptors are only deflected by BC vibration.

| HEARING OWN VOICE: Routes to each ear |  |  |  |  |
| --- | --- | --- | --- | --- |
| Route | Sensory organ | Arrival time after vocalisation | Interaural arrival time and intensity | Comment |
| BC | Vestibular system | ~ 0.5 ms | Identical (assumes body symmetry) | Insensitive to environmental variation. BC attenuation and filtering are consistent in the short- and medium-term, with only small and gradual long-term changes which follow head composition and body profile across the lifespan. |
| BC | Cochlea | ~ 0.5 ms |  |  |

**Table 1: Sound and vibrational energy is transmitted to each ear through body conduction (BC) and direct and reflected air conduction (dAC/rAC), and can deflect two sets of mechanoreceptors in each inner ear.**
| **HEARING OWN VOICE: Routes to each ear** |  |  |  |  |
| --- | --- | --- | --- | --- |
| Route | Sensory organ | Arrival time after vocalisation | Interaural arrival time and intensity | Comment |
| dAC | Cochlea | ~ 0.5 ms | Near identical (can vary with head orientation and air turbulence) | During vocalisation, dAC and BC contributions are approximately equal at the cochlea (von Békésy, 1949; Pörschmann, 2000; Reinfeldt et al., 2010). Some spectral variation (e.g. nasal phonemes more prominent over BC) due to filtering differences between air and body. Recordings of own speech (capturing predominantly dAC) are often found by the speaker to differ from what is heard while speaking (a mixture of BC and dAC, with some rAC). |
| rAC | Cochlea | typically 2–50 ms | Different | Arrives within 2–50 ms (or longer, depending on environment) of dAC sound and BC vibration. Less sonic/vibrational energy than the dAC/BC mixture. Delay relative to dAC/BC creates comb filtering. Delays of rAC above ~50 ms are experienced psychoacoustically as an echo; delays of rAC below ~50 ms are psychoacoustically fused with dAC/BC as in the Haas or precedence effect. |

Groups of neurons having response properties supporting coincidence detection on the millisecond timescales required for the hypothesised own voice identification mechanism can be found in the cochlear nucleus and superior olivary complex. Review of brainstem neurons can be found in Golding & Oertel (2012) and review of vestibular inputs to the cochlear nucleus in Newlands et al. (2003) or Smith (2012). Figure 3 shows a sagittal view of brain areas innervated by the inner ear, and figure 4 shows cortical areas with connectivity to the vestibular system alongside areas important for speech and language. The cochlear nucleus and superior olivary complex comprise initial stages in a subcortical chain referred to as the ascending auditory pathway (Irvine, 1992). Changes in firing rates within brainstem neurons which correspond to the hypothesised coincidence detection could in turn be expected to change activity at higher stages of the ascending auditory pathway, including inputs to the cortex. Such activity could be interpreted as an auditory stream which identifies own voice.

**Figure 3:**
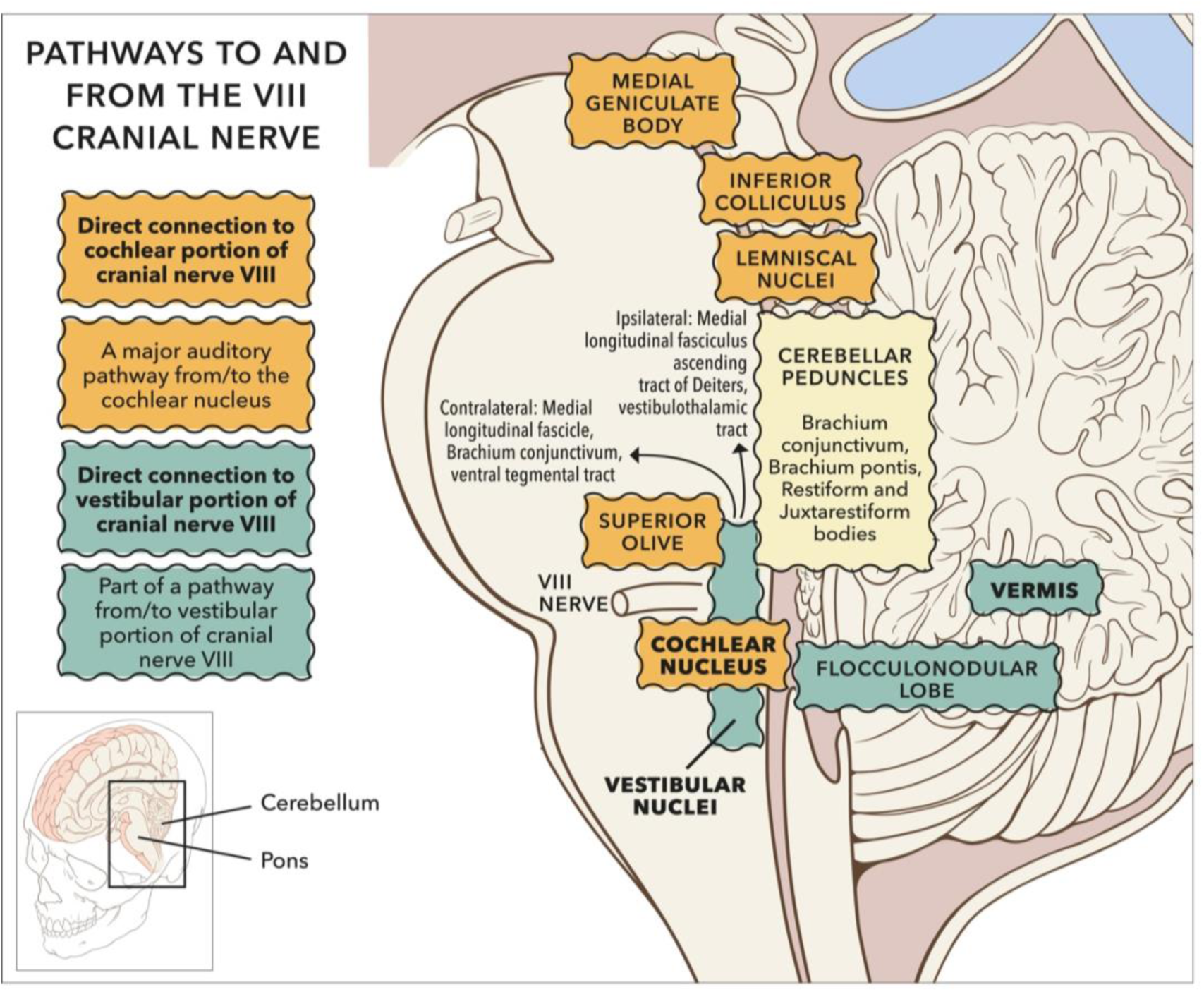
Sagittal view of subcortical pathways to and from the VIII cranial nerve. Whilst the auditory pathway ascending from the cochlear nucleus is relatively well established (Irvine, 1992), pathways to and from vestibular nuclei remain under investigation (Pierrot-Deseilligny & Tilikete, 2008; Zwergal et al., 2009). Investigation is largely using animal models. Projections to vestibular cortex via the thalamus have been established in humans through clinical observation and lesion studies (Conrad et al., 2014; Hitier et al., 2014; Wijesinghe et al., 2015). Vestibular nuclei also project down the spine (not shown). © Portions of this figure were adapted from illustrations by Patrick J. Lynch, http://patricklynch.net/. Creative Commons 2.5 licence.

**Figure 4:**
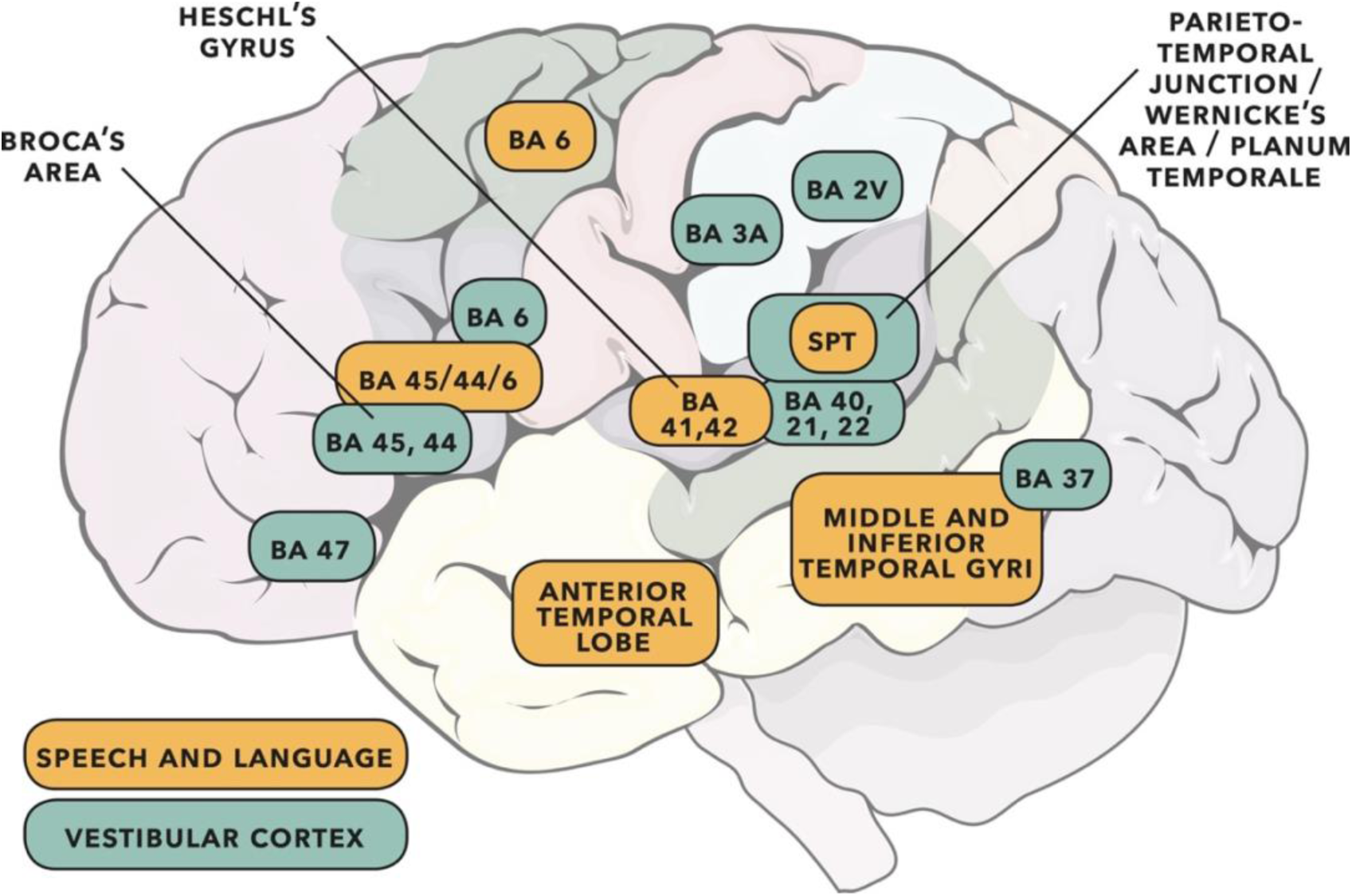
Cortical areas important for speech and language (adapted from the dual-stream model of Hickok & Poeppel, 2007) shown with vestibular cortical areas identified in cats, monkeys and humans (adapted from Ventre-Dominey, 2014; see also Frank & Greenlee, 2018). Cortical activity following vestibular input has wide interpretation (e.g. see reviews of cognition in Hitier et al., 2014, and auditory/rhythm/timing in Todd & Lee, 2015). Some of the vestibular areas identified will be predominantly related to gravitoinertial function (see discussion in Ferrè & Haggard, 2020). Numbers are Brodmann areas – see primary literature for more exact location detail. Spt is the Sylvian parieto-temporal region proposed by Hickok & Poeppel (2007) as a sensorimotor integration area. Vestibular sites in humans have been identified as such when direct electrical stimulation of the cortex gives rise to gravitoinertial illusion. When vestibular sites are identified within BA 21 (lateral temporal lobe) or BA 22 (Wernicke’s area), auditory illusion is found to accompany gravitoinertial illusion (Kahane et al., 2003; Fenoy et al., 2006). © Portions of this illustration were adapted from Servier Medical Art, https://smart.servier.com. Creative Commons 3.0 licence.

##### 2.3.2.1 Explanatory Power

As a process explanation, the Concurrency Hypothesis provides a detailed account of how own voice is identified. The proposed involvement of particular types of brainstem neurons (e.g. octopus cells in the cochlear nucleus, or bipolar principal cells of the medial superior olive) generates testable auxiliary hypotheses (see discussion in sections 2.4 and 3.4). Whereas the existence of an own voice auditory stream, which is identified through coincidence detection between vestibular and cochlear afferents, is the core hypothesis.

There is also a contrastive explanation of why own voice is identified in the way described by the Concurrency Hypothesis. The contrastive explanation addresses evolutionary and philosophical considerations. The Concurrency Hypothesis as described so far is specific to mammals. However, the Concurrency Hypothesis could be extended to all terrestrial and amphibious vertebrates if the basilar papilla is considered in place of the cochlea; to fish if the lagena is considered; and in principle to any animal which produces sound and vibration, and has two or more sets of sensory receptors capable of detecting sound and vibration. See species surveys in Suthers, Tecumseh Fitch, Fay & Popper (2016) and Pollack, Mason, Popper & Fay (2019).

The prospect of such a wide taxonomic application for the Concurrency Hypothesis suggests a provenance early in evolution. This in turn prompts reconsideration of the role of the inner ear. The Concurrency Hypothesis provides a principled distinction between self (identification of own voice) and environment (reflection of own voice from surroundings). Such a distinction has importance for cognitive science and philosophy of mind (Wilson & Foglia, 2017). For example, in a representational theory of mind the distinction between self and environment is integral to content determination (Pitt, 2020).

The basis for the self-environment distinction in the Concurrency Hypothesis is the presence of two sets of mechanoreceptors in the ear. One set of mechanoreceptors detects own voice in isolation, the other detects own voice mixed with ambient sound, including reflection of own voice. This is dissimilar to other modalities. For example, the visual analogy would be identification of one’s own hand. However, photoreceptors do not collect sufficient information to identify one’s own hand from light waves incident on the retina. Such identification would be possible following multisensory integration, but this is also the case for audition (e.g. as in the combination of audition with proprioception during vocalisation).

As such, audition might be the only modality within which self and environment can be distinguished. If so, multisensory integrations including audition could underlie self-environment distinction for modalities other than audition. Evolution of any such dependency would have to create phenotypes sufficiently robust to account for self-environment distinction when hearing ability is absent. Further consideration of such matters is beyond the scope of this article, but would follow discussions of heritability and innateness such as those in Griffiths (2020), Godfrey-Smith & Sterelny (2016) or Downes & Matthews (2020).

Self-environment distinction is also important in our understanding of consciousness (Van Gulick, 2018). For example, our experience of qualia depends on introspection from what we presume to be a shared environment. Our intentionality towards objects other than ourselves rests likewise. From considerations such as these, provision of a principled basis for distinction between self and environment would be a comparably important function of the inner ear as its hearing function.

### 2.4 Discussion

This section describes a general application of the Concurrency Hypothesis to speech-motor research and auditory scene analysis. Section 3 will build on the discussion in this section to describe a specific application of the Concurrency Hypothesis to explanation of stuttering.

#### 2.4.1 Application to speech-motor research

An own voice auditory stream would provide a target for the proposed efference copy of the speech plan in predictive feedback control models (e.g. Hickok & Poeppel, 2007; Roelofs & Meyer, 1999; Tourville & Guenther, 2011). If applied to speech-motor models, the Concurrency Hypothesis has potential to improve explanatory power.

A corollary of this proposal is that if the Concurrency Hypothesis is to be tested, speech-motor research should use physiologically valid own voice stimuli. Physiologically valid own voice stimuli are those containing concurrent AC sound and BC vibration, with relative composition and timing as described in figure 1 and table 1.

Creation of such stimuli carries practical difficulty. For example, an ideal test of speech-motor activity would compare brain activity during identical sound and vibrational stimuli in two conditions. The first condition is the standard articulatory process: brain activity generates sound and vibration following coordinated nerve impulses to articulatory muscles, whilst at the same time brain activity is altered following deflection of inner ear mechanoreceptors by the sound and vibration produced during articulation. The second condition should be identical to the first, but without the activity in articulatory muscles being created by brain activity. Instead, the measured brain activity would be solely in response to the sound and vibration produced by articulatory muscles. Unfortunately, the experimental arrangement in the second condition is difficult or impossible even in animal models. The articulatory muscles could in principle be made to produce sound and vibrational stimuli similar to that during vocalisation, for example through electrical stimulus to the articulatory muscles. However, the process of doing so would either be highly traumatic to the host animal, or the animal would have to be sedated. Whatever experimental arrangement is chosen, resting state brain activity in the second condition would differ from that of the first condition (the standard articulatory process) to the extent that comparison of brain activity between the two conditions would be overwhelmingly difficult to interpret.

Accordingly, much testing of brain activity during articulation, or vocalisation, has been based around a simpler comparison. The first condition is the standard articulatory process (i.e. as previously defined), with simultaneous recording of brain activity (e.g. by electrophysiology) and the sound and/or vibrations created during articulation (e.g. using a microphone). The second condition comprises a recording of brain activity without articulation, whilst the sound and/or vibration recorded in the first condition is played back. This comparison would seem to overcome the difficulty with having articulatory muscles create the sound and vibration in the second condition. However, there is a disanalogy in that the sound and vibration in the second condition are not identical to the sound and vibration in the first condition. This disanalogy has potential to invalidate the intended comparison.

Thus, protocols intended to compare brain activity during articulation and the playback of a recording of vocalisation must choose a methodology for recording and playback of the sound and/or vibration. Possibilities are shown in a Latin square in figure 5. Of these, speech-motor investigation has overwhelmingly compared the own voice condition with dAC playback of a dAC recording. Often, participants are invited to adjust sound pressure levels of dAC playback so as to perceptually match the loudness of the AC/BC combination heard during vocalisation. Doing so does not create a stimulus comparable to the stimulus present during vocalisation. Own voice is perceived through an approximately equal combination of air- and body-conducted stimuli (von Békésy, 1949; Pörschmann, 2000; Reinfeldt et al., 2010). Perceptual doubling of the loudness of the AC stimulus, to compensate for the absence of BC stimulus, will for most participants correspond to no more than a 10 dB increase in sound pressure level (Stevens, 1972; Warren, 1973; Florentine, Popper & Fay, 2010). Such an increase will barely bring the AC stimulus to vestibular threshold, which for AC is 10 dB above the 60 dBA level typical of conversational speech. The AC vestibular threshold is moreover 35 dB above the BC vestibular threshold (McNerney & Burkard, 2011; Welgampola, Rosengren, Halmagyi & Colebatch, 2003).

**Figure 5:**
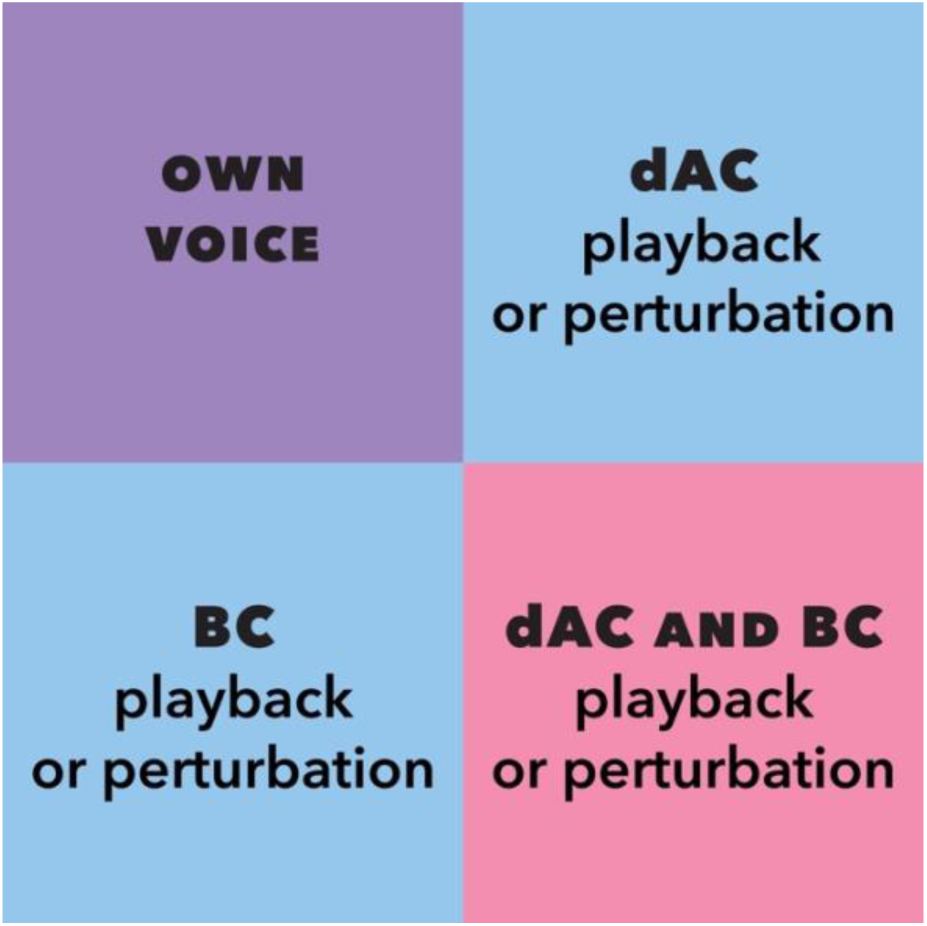
Latin square showing sound and vibrational stimuli which could be used in brain studies of own voice audition. The “own voice” condition is the standard articulatory process. It includes speech-motor brain activity which results in articulation generating dAC and rAC sound, and BC vibration; and at the same time includes the brain activity following deflection of inner ear mechanoreceptors by the dAC and rAC sound, and BC vibration, produced during articulation. “Playback” refers to playback of recordings of sound or vibration made during the standard articulatory process. Playback conditions do not contain speech-motor activity, unless digitally processed playback with a short delay (usually 10 ms or more) is presented concurrently with ongoing articulation. Such short latency digital manipulation is referred to as perturbation, and may also include manipulations to recordings (e.g. frequency shifts or changes to the nature of formants). A limitation for any type of playback is that the sound and vibrational stimuli present in the own voice condition cannot be recreated exactly using the earphones and bone vibrators available in laboratories. Combined air- and body-conducted (AC/BC) playback according to the timings provided in table 1 (i.e. AC and BC playback with binaural arrival at the inner ear coincident to ∼ 0.1 ms) offers the closest approximation to the sound and vibrational stimuli present in the own voice condition. Not shown in the diagram is that BC stimulus can be subdivided into levels above and below vestibular threshold. BC stimulus should be above vestibular threshold, and AC stimulus below vestibular threshold, to mimic stimuli present during articulation.

It follows that even after a sound pressure level increase to perceptually match the loudness of own voice, stimulation due to AC playback will either deflect vestibular mechanoreceptors very weakly in comparison to the BC stimulation present during vocalisation, or stimulation due to AC playback will not deflect vestibular mechanoreceptors at all. Firing rates of the vestibular ganglion will be altered barely or not at all from resting state. Action potentials along the VIII cranial nerve will predominantly be altered according to deflection of cochlear mechanoreceptors by AC playback, and an auditory stream corresponding to own voice will not be identified through coincidence detection between cochlear and vestibular streams as per the Concurrency Hypothesis.

Many functional imaging studies have compared vocalisation to AC playback of own voice recordings (e.g. with human participants: Numminen et al., 1998; Numminen & Curio, 1999; Curio et al., 2000; Ford et al., 2001; Houde et al., 2002; Ford & Mathalon, 2004; Ventura et al., 2009; Greenlee et al., 2011; Sato & Shiller, 2018; or using animal models: Müller-Preuss & Ploog, 1981; Eliades & Wang, 2017; Eliades & Tsunada, 2018). A consistent finding in such experiments is that parts of temporal cortex which respond to sound have reduced activity in the vocalisation condition compared to the playback condition. This has been interpreted as speech-motor activity modulating the temporal cortex (Hickok et al., 2011; Parrell & Houde, 2019). The interpretation is consistent with theoretical models in which attenuating auditory feedback increases accuracy of state estimates of the speech-motor system (Parrell et al., 2019).

Whilst an attractive explanation, motor induced suppression of temporal cortex is not strongly supported by studies comparing vocalisation and AC playback conditions. The reason for this is that vocalisation and playback stimuli differ (as per figure 5), meaning that the observed reduction in temporal cortex activity cannot conclusively be attributed to speech-motor activity modulating temporal cortex. An alternative explanation is that the observed reduction in temporal cortex activity is due to the difference in stimuli between vocalisation and AC playback conditions. The Concurrency Hypothesis is consistent with this alternative explanation. The Concurrency Hypothesis adds the detail that in the vocalisation condition, firing rates of neurons in the ascending auditory pathway will uniquely identify own voice through coincidence detection of cochlear and vestibular afferents. Whereas in the AC playback condition, the ascending auditory pathway functions as it would with any ambient AC stimulus (i.e. as per Irvine 1992; Bregman, 1990).

It is possible that both explanations are correct: that an own voice auditory stream modifies temporal cortex activity, and that articulation modifies temporal cortex activity independently of audition. Exploring these possibilities offers the opportunity to increase explanatory power of speech-motor models, and to make testable predictions. In doing so it is not necessary to use the Concurrency Hypothesis. However, alternatives would be to propose a different method by which own voice is identified as an ascending auditory stream (i.e. a solution to the ill-posed problem of sound source discrimination in auditory scene analysis), or else to stipulate that an auditory target map for own speech is innately specified (e.g. as per Liberman & Mattingly, 1985).

Studies using playback of own voice recordings could be reinterpreted in light of these considerations, and extended to include BC stimuli. Auditory perturbation studies could be similarly reinterpreted (e.g. McGuire et al., 1996; Hirano et al., 1997; Fu et al., 2006; Parkinson et al., 2012; Toyomura et al., 2007; Zarate & Zatorre, 2008; Tourville et al., 2008; Zheng et al., 2009; Zarate et al., 2010). In auditory perturbation studies, vocalisation is recorded, is optionally digitally manipulated, and is played back with a short delay whilst articulation is ongoing. Examples of manipulation include frequency shift or alteration of formants. Recording and playback use AC sound. Digital processing (e.g. with fast Fourier transform) introduces delays which are typically 10 ms or more. Such delays are at least an order of magnitude larger than the sub-millisecond timings in table 1. Thus, auditory perturbation studies assess the effect of keeping the BC vibrational stimulus of vocalisation unchanged, whilst adding a delayed AC stimulus having similar spectral characteristics to the ongoing vocalisation. Effectively they manipulate rAC and (if using insert earphones) attenuate dAC. The protocol could be extended to form part of a larger range of investigation in which BC, and combined AC/BC, manipulations are also evaluated.

The Latin square in figure 5 is a simplification. Stimuli can be further subdivided into those above and below vestibular threshold. Todd et al. (2014a, 2014b) compared cortical response to stimuli above and below vestibular threshold. Electroencephalography showed morphological change in and around the N1 wave upon crossing vestibular threshold, with source analysis indicating origin in cingulate or temporal cortex. The N1 wave (or its M100 equivalent in magnetoencephalography) is the component found to have reduced amplitude when brain activity during vocalisation is compared to brain activity during AC playback of vocalisation. Thus, the suggestion is that in studies comparing vocalisation and playback conditions, the observed brain activity will differ depending on whether playback stimuli are above or below vestibular threshold. A physiologically valid own voice stimulus will combine BC stimulus above vestibular threshold with AC stimulus below vestibular threshold. Follow-up work to the current article will appraise brain activity following combinations of BC and AC stimuli which are respectively above and below vestibular threshold.

#### 2.4.2 Application to Auditory Scene Analysis

Bregman (1990) proposed that auditory scenes are generated from the neural firing patterns elicited when sound waves are coincident on the biomechanical structure of the middle and inner ears. Auditory scenes would contain detail consistent with our perceptual experience. Two processes are proposed to identify the auditory streams which comprise auditory scenes. Firstly, primitives, which are general purpose segregation and grouping processes based on those developed by the Gestalt school (e.g. common onset, harmonicity, spectral composition, co-variation in amplitude; Carlyon, 2004; Darwin, 2007; Ciocca, 2008; Denham & Winkler, 2015; Młynarski & McDermott, 2019). Secondly, schemas, which are specific processes identifying certain types of sound (e.g. conspecific animal vocalisations or phonemes in human speech; Bey & McAdams, 2002; Billig et al., 2013; Woods & McDermott, 2018).

The Concurrency Hypothesis could be the basis of a schema identifying own voice. Modelling of auditory scene analysis is an active research area (Cooke & Ellis, 2001; Haykin & Chen, 2005; Snyder & Alain, 2007; Winkler et al., 2009; Szabó, Denham & Winkler. 2016; Snyder & Elhilali, 2017; Chakrabarty & Elhilali, 2019). Whichever modelling approach is taken, the Concurrency Hypothesis would be applied through the following principles:

i. Primitive processes are proposed to act on neural firing patterns elicited by deflection of vestibular mechanoreceptors as well as by deflection of cochlear mechanoreceptors.
ii. Whenever firing patterns of vestibular and cochlear origin have similar attributes as identified by primitives, the firing patterns are likely to correspond to own voice.
iii. Activity in the auditory brainstem (BinKhamis et al., 2019) is consistent with substantial processing of speech sounds. As such, models will have greater neurological plausibility if the coincidence detection in (ii) occurs very early in the ascending auditory pathway – for example, in the cochlear nucleus or the superior olivary complex.
iv. Computational modelling of coincidence detection (e.g. through vestibular input to octopus cells in the cochlear nucleus) may require primitives and schemas to be entwined. An own voice identification schema based on (i – iv) could underpin further schemas. Possibilities are:
v. Vocalisation of conspecifics is likely to be occurring when primitives identify similar neural firing patterns (e.g. spectral composition typical of formants) to those present during own voice coincidence detection, but when vocalisation is not being produced and neural firing patterns arise from cochlear mechanoreceptors only.
vi. If stored in short-term memory, an own voice auditory stream could be compared via primitives to the rAC reflections of own voice (see figure 2 and table 1) to create a schema identifying reflection and reverberation.
vii. Multisensory integration (Stein & Stanford, 2008) of reflections and reverberations from (vi) with head and body position could support a schema for echolocation (see review of human echolocation in Kolarik et al., 2014).
viii. Sound source learning based on (vii), in combination with the generalised vocalisation schema of (v), could support a schema distinguishing sources in multi-speaker scenarios.
ix. Adaptation of the schema in (viii) for sounds other than vocalisation could reinforce learning of sound source location using primitives.

These ideas need development into computational models. The underlying point is that many or all of the schemas required by auditory scene analysis could be based on the Concurrency Hypothesis. The high energy vocalisations of neonates (e.g. crying or wailing) have more than sufficient energy to deflect both cochlear and vestibular mechanoreceptors, meaning that auditory learning based on the Concurrency Hypothesis would begin at birth (and quite possibly, would have a precursor based on the mother’s voice in utero).

## 3. Hypotheses of Stuttering

### 3.1 Explanatory targets

Explanatory targets for stuttering are extensive. Table 2 shows process explananda (how stuttering happens), whilst table 3 shows contrastive explananda (why stuttering happens). These lists are not intended as exhaustive, but are rather presented as minimal criteria which any hypothesis of stuttering should address.

**Table 2:**
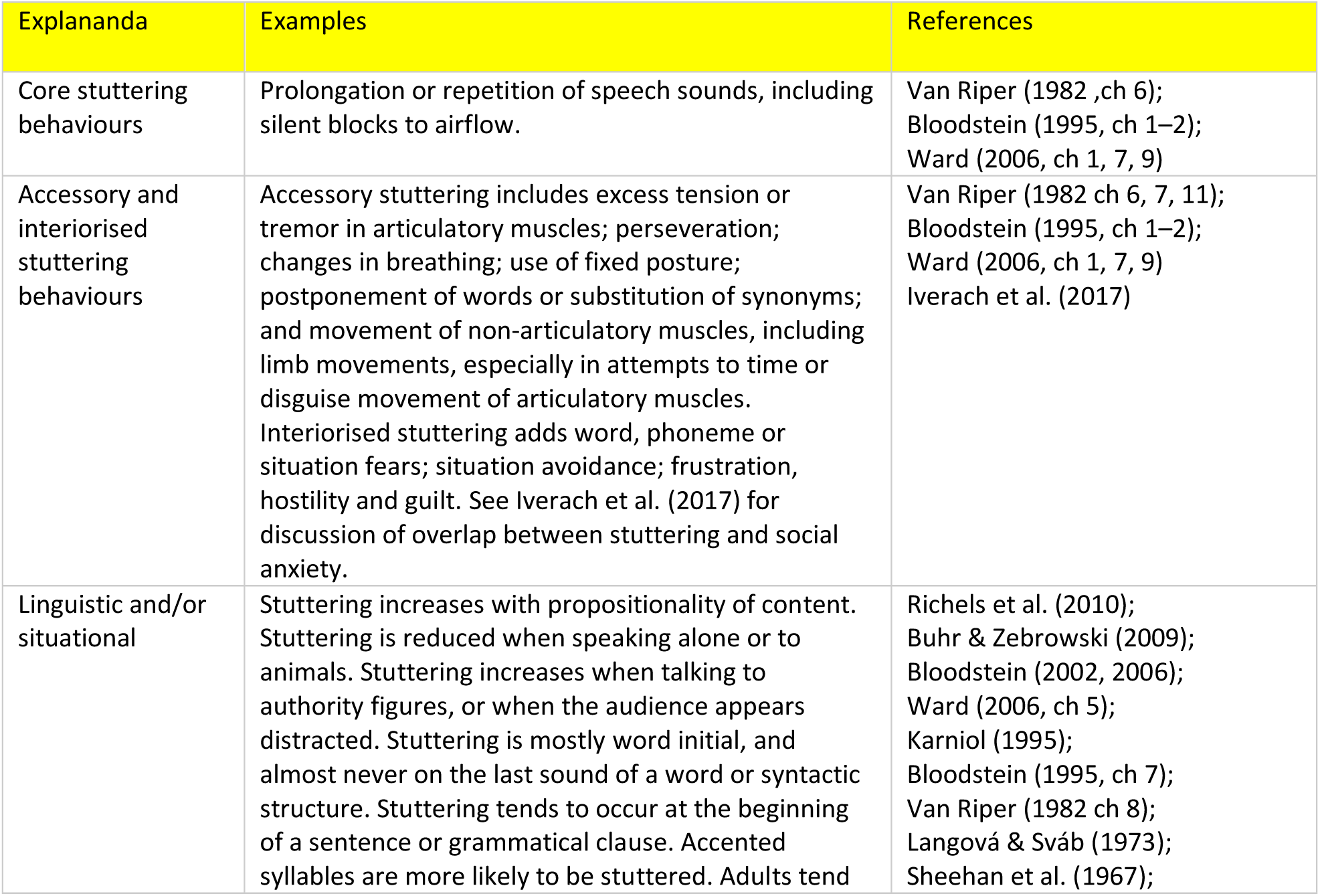
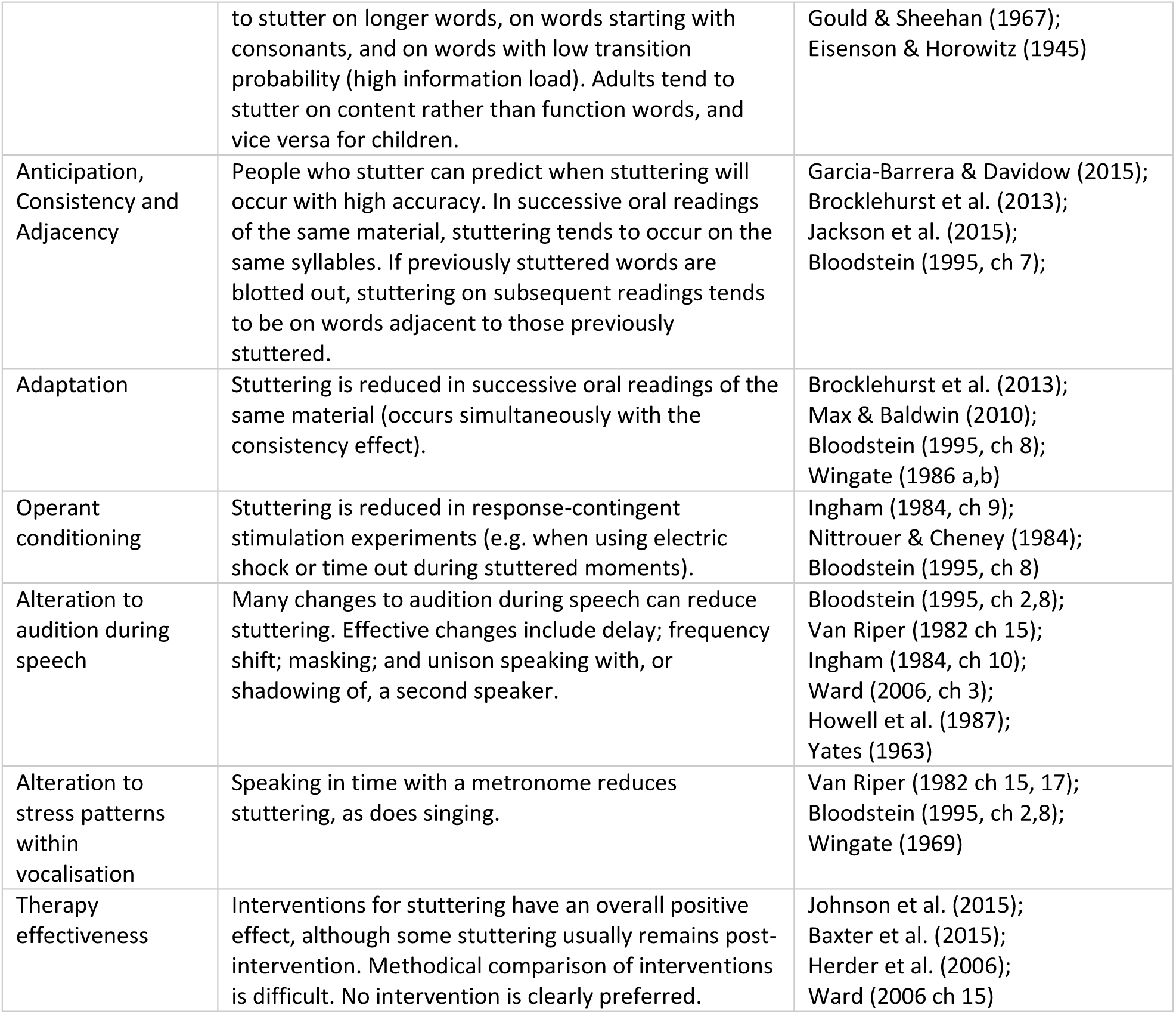
Process explananda for stuttering

**Table 3:**
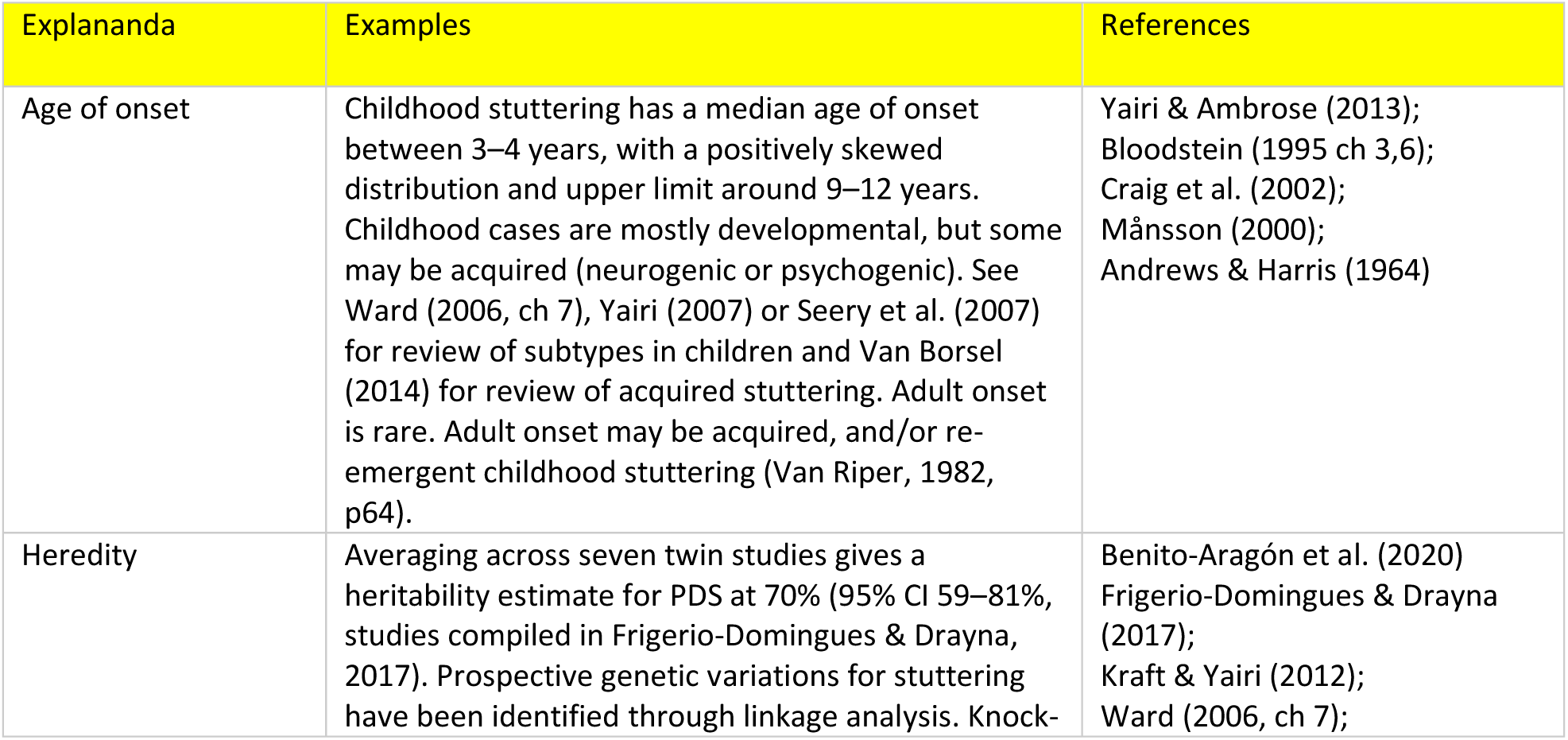
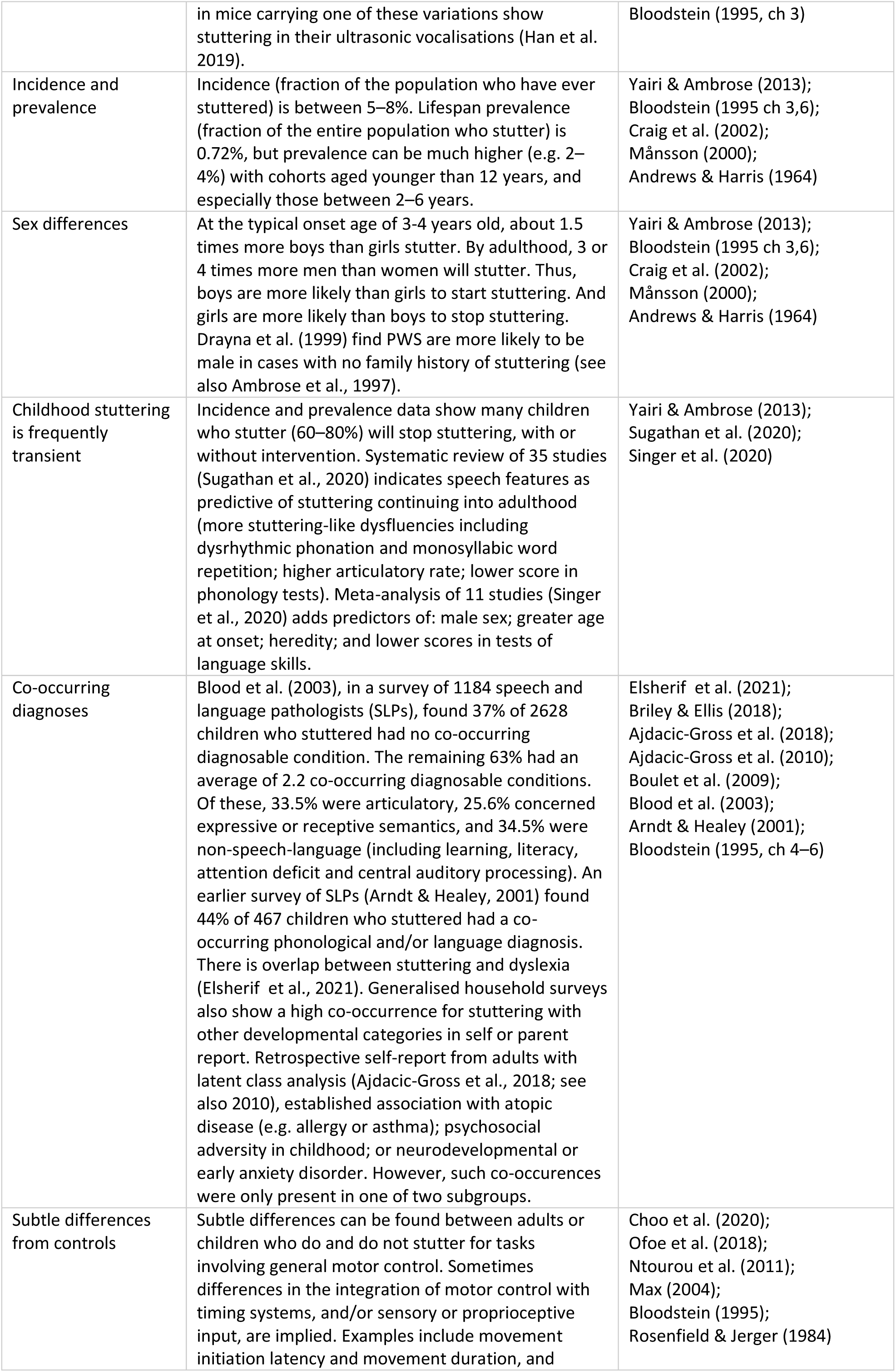
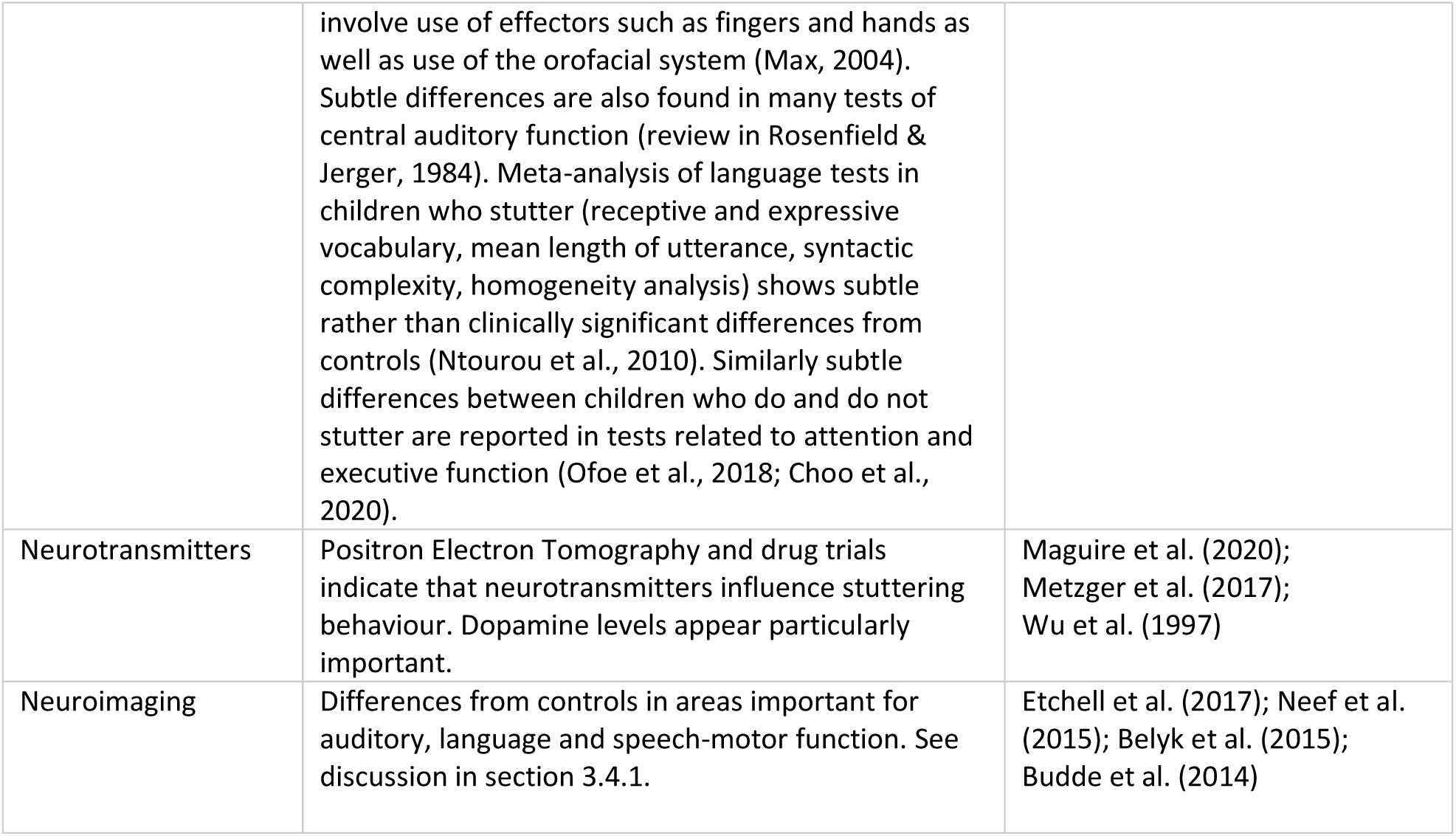
Contrastive explananda for stuttering

Priority will be given to addressing process explananda. This is not to downplay the importance of contrastive explananda for stuttering research. However, a comprehensive discussion of contrastive explananda for stuttering (e.g. why there is a sex difference; the role of heredity; whether a particular brain study reflects causation, consequences or correlates of stuttering) encompasses issues wider than those within stuttering research, and is accordingly outside the scope of this article. The current aim is of adequacy for process explanans, with contrastive explanans added as part of ongoing research.

### 3.2 Candidate explanations

Bloodstein (1995) categorises hypotheses of the moment of stuttering into three groups: repressed needs, anticipatory struggle, and breakdown. Research and theoretical development over the last 30 years has overwhelmingly focussed on breakdown hypotheses. As such, repressed needs hypotheses and anticipatory struggle hypotheses will be reviewed only in brief, whilst breakdown hypotheses will be described in greater detail.

#### 3.2.1 Repressed Needs Hypotheses

Originating in the psychoanalytic schools of the 1920s and 1930s, repressed needs hypotheses describe stuttering as a neurotic symptom rooted in unconscious needs. Such hypotheses are outside the mainstream of contemporary stuttering research (Martin, 2016).

#### 3.2.2 Anticipatory Struggle Hypotheses

In anticipatory struggle hypotheses, stuttering is preceded by the speaker’s prediction that speech will be difficult to execute. The prediction of difficulty leads to increased muscular tension. The increased muscular tension in turn impairs the coordination usually present during speech, and causes the speech attempt to be stuttered.

Anticipatory struggle hypotheses have seen little development in the last 50 years. For a historical survey, see Bloodstein (1995, ch 2), and for a contemporary perspective see Brocklehurst et al. (2013).

#### 3.2.3 Breakdown Hypotheses

In breakdown hypotheses, stuttering is a behavioural manifestation of vulnerability in speaking ability. The vulnerability is generally proposed to occur in either the language encoding or the speech-motor system. Breakdown of the vulnerable system is typically attributed to emotional or psychosocial stress (Bloodstein, 1995, p60).

##### 3.2.3.1 Language encoding breakdown

Language encoding breakdown has been described in what Levelt (1989, 1999) refers to as the Formulator. This is a hypothesised stage of speech production between thought and expression, in which lexical and syntactic selection, along with morphological, phonological and phonetic encoding, precedes creation of a motor plan. Levelt’s model is shown in figure 6. The Formulator can be described using a spreading activation network (e.g. Dell, 1986; Dell & O’Seaghdha, 1991). In network models, a metrical frame is created for a planned utterance. Phonological segment nodes will then compete for selection, with the nodes filling the frame being those which have the highest activation level at the moment when speech-motor planning commences.

**Figure 6:**
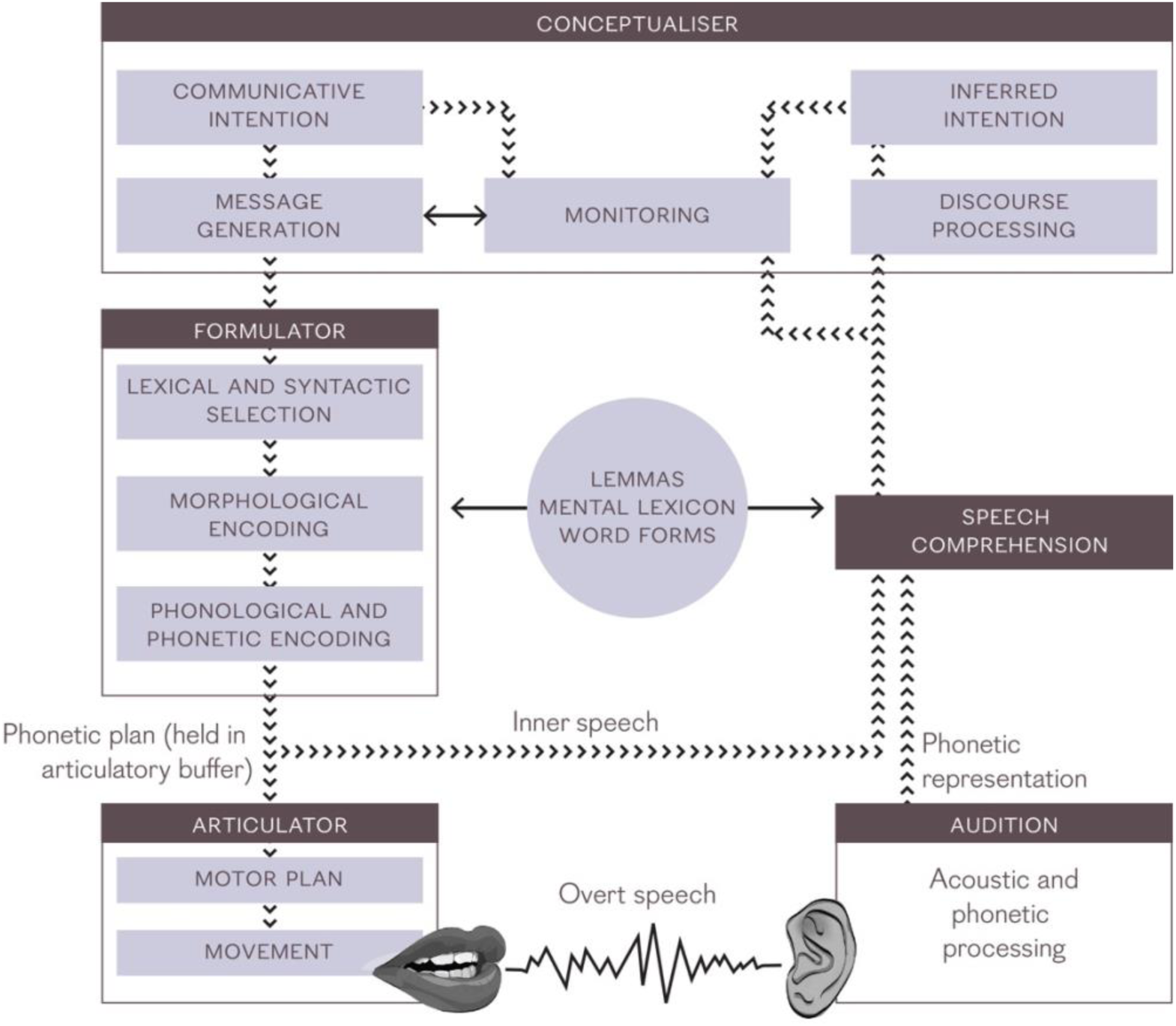
Speech production model of Levelt (1989; see also Levelt et al., 1999). Notable features are an inner and an outer loop, with the parsimony of a shared mental lexicon. Stages include Audition, Speech Comprehension, Conceptualisation, Formulation and Articulation. The Concurrency Hypothesis concerns activity in Audition, and thus addresses a special case of auditory scene analysis (Bregman, 1990). Discussion of Speech Comprehension can be found in Norris et al. (2000), Galantucci et al. (2006) or Poeppel et al. (2008), among others. There is no widely agreed model of the Conceptualiser; any effort to produce one touches on long-standing issues in Cognitive Science, Philosophy of Psychology and Philosophy of Mind (several other hypotheses of the Levelt model, and hypotheses of its constituents, do likewise). Indefrey & Levelt (2004) present a meta-analysis, based on neuroimaging literature, of the time course for processes within the Formulator; see also section 3.2.3.1 for discussion of Dell’s (1986) spreading activation network model of the Formulator. Articulation is described by speech-motor control models such as DIVA (Guenther et al., 2006) or FACTS (Parrell et al., 2019). © Creative Commons 4.0 licence. Based on Levelt (1989), Levelt et al. (1999), Indefrey & Levelt (2004).

The Covert Repair Hypothesis (Postma & Kolk, 1993) postulates slower than usual activity in the Formulator for PWS. As a result, a speech plan may be created whilst nodes are still competing for selection. If inappropriate nodes are selected, two possibilities pertain. If the inappropriate nodes are detected prior to articulation (e.g. via an internal monitoring loop), they are repaired covertly. This repair manifests as a silent pause – the speaker wishes to continue, but cannot do so at that moment.

Alternatively, if inappropriate nodes are detected during articulation, the speaker will stop and retrace. Phonemes uttered prior to retrace are audible as stuttering for however many reformulations are necessary to correct the speech plan. A variant on this theme is offered by the Vicious Circle Hypothesis (Vasić & Wijnen, 2005; Bernstein Ratner & Wijnen, 2007), which proposes that it is over-vigilance in repair, rather than slower than usual formulation, which causes stuttering.

An alternative breakdown mechanism is described by Howell (2004, 2008). In the EXPLAN hypothesis, breakdown occurs when the rate of speech planning has fallen below that of execution. The available speech plan is repeatedly executed until a continuation of the speech plan is available. EXPLAN entails aspects of both psycholinguistic and speech-motor breakdown. The Variable Release Threshold hypothesis (Brocklehurst et al., 2013) modifies EXPLAN such that the release threshold for a phoneme will vary according to a modified version of Bloodstein’s (1975) account of anticipatory struggle.

##### 3.2.3.2 Speech-motor breakdown

Speech-motor breakdown is typically investigated through comparison of people who stutter in fluent versus stuttered speech (state comparison) or people who do and do not stutter during fluent speech (trait comparison). Outcome measurement is via neuroimaging, electromyography of articulatory muscles, or a hybrid design (e.g. studies employing transcranial magnetic stimulation). Differences are reliably and repeatedly established in both trait and state comparisons, and are present even below the threshold for behavioural observation of stuttered speech (Etchell et al., 2017; Neef et al., 2015; Belyk et al., 2015; Budde et al., 2014). Brain areas frequently identified include premotor cortex and the temporo-parietal junction (including white matter connecting those areas), the cerebellum, and the basal ganglia. Stuttering can be emulated neurocomputationally by modelling the brain activity observed in neuroimaging of stuttering (Civier et al., 2013), with over-reliance on auditory feedback a contributing factor (Max et al., 2004; Civier et al., 2010). Arenas (2017) proposes an extension to speech-motor breakdown in which fluctuations in the vigilance of the monitoring system account for the contextual variability of stuttering.

### 3.3 A Novel Account of Stuttering: REMATCH (Reflexivity and Communicative Mismatch)

#### 3.3.1 Introduction

This section introduces a novel account of stuttering with two core hypotheses: Reflexivity, and Communicative Mismatch. The combination is referred to as REMATCH.

The first core hypothesis in REMATCH concerns a quale referred to as “reflexivity”. It proposes that PWS have a subjective experience during speaking in which their own speech has increased salience in comparison to the way that people who do not stutter experience their own speech while speaking. The second describes communicative mismatch, in which a breakdown in communicative choreography between speaker and listener engenders observable stuttering behaviour. The reflexivity proposal develops the Concurrency Hypothesis described in Section 2. It is a distal cause of stuttering relative to communicative mismatch.

This section will proceed as follows. The sequence of events leading to a moment of stuttering, consistent with the two core hypotheses, will be described. The core hypotheses will then be applied to the explananda in tables 2 and 3.

#### 3.3.2 Increased Reflexivity

Consider that the subjective experience of seeing the colour red may differ between individuals, even if those individuals can mutually agree that the referenced colour is red (Tye, 2018). Similarly, different speakers may have differing subjective experiences of hearing their own voice during vocalisation. The proposal is that the subjective experience of hearing own voice during vocalisation differs in a principled and consistent manner between people who do and do not stutter.

This subjective experience, or quale, of own voice during vocalisation will henceforth be referred to as “reflexivity”. It is related to self-awareness (Gallagher & Zahavi, 2021; Smith, 2020). The exact proposal is that reflexivity is increased for PWS relative to controls. What is meant by increased reflexivity is that the phenomenal experience of own voice is more intense for PWS than for ordinarily fluent speakers. It is as if PWS were speaking through a magical megaphone, which broadcasts only inside the body, and whose effect is to increase salience of the message being delivered rather than volume of the utterance.

Empirical investigation of qualia is achievable through psychophysics, albeit with well-identified difficulties (Fodor, 1987). The proposal of reflexivity as a quale builds on the Concurrency Hypothesis described in Section 2, and in particular it follows from the issues around evolution, cognitive science and philosophy of mind discussed in section 2.3.2.1. The hypothesis of Communicative Mismatch, to be introduced in section 3.3.1.2, proposes that a difference in subjective experience of the reflexivity quale between people who do and do not stutter is causative of stuttering behaviour.

A difference in reflexivity between people who do and do not stutter could be expected to coincide with a difference in the auditory feedback whose presence is integral to many types of psycholinguistic and speech-motor models. Alterations to auditory feedback are well-established as reducing stuttering for PWS (Yates, 1963; Howell et al., 1987; Kalinowski et al., 1993; Stuart et al., 2004; Foundas et al., 2013), and hyperfunctional monitoring in stuttering has been proposed from psycholinguistic (Bernstein Ratner, 1997; Bernstein Ratner & Wijnen, 2007) and speech-motor (Arenas, 2017) perspectives. REMATCH is independent of any particular speech-motor or psycholinguistic model. For example, speech may be entirely under feed forward control, or else speech may be best described by paradigms which do away with mental representation entirely (e.g. certain types of dynamical system, or those of extended cognition). For readers who prefer to think in terms of feedback control, the idea would be that an entire person (including the history, personality, hopes, dreams, and so forth) is included in the feedback loop for own voice audition. See Mysak (1969, ch 7) for a systems control account of stuttering along these lines.

#### 3.3.3 Communicative Mismatch

When PWS describe moments of stuttering, the role of the audience and situation are among themes identified (Tichenor & Yaruss, 2018). In a review of linguistic factors, Karniol (1995) suggests that the involvement of motor process in stuttering is a symptom rather than a cause. Pierre (2015), extending beyond linguistics to discuss societal convention more broadly, describes stuttered speech as marginalised relative to dominant choreographies of bodily and inter-bodily communicative practices.

To address perspectives such as these, stuttering will be considered not just as an interruption of speech, but moreover as an interruption of a speech act (i.e. speech act as per Austin, 1955). It will be based around the approach-avoidance conflict hypothesis of stuttering developed by Sheehan (1953, 1958, 1970, 1975). Approach-avoidance conflict was originally formulated as a Gestalt field theory by Lewin (1935). Conflict would follow incompatible goals – for example, accept a substantial pay rise (approach gradient), but only with unpaid weekend work at the employer’s discretion (avoidance gradient). Sheehan (1958) proposed that stuttering is a double approach-avoidance conflict, in which “[The person who stutters] can speak, thus achieving his aim of communication, but at the cost of the shame and guilt he has learned to attach to his stuttering. Or he can remain silent, abandon communication, and suffer the frustration and guilt that such a retreat carries with it.”

Sheehan was inspired by the work of Miller (1944) who trained rats in a runway first with a food goal, then with electric shock. When Miller presented the previously trained rats with a combination of a food goal and electric shock, the rats would display motor control vacillations similar to those observed in stuttering. In an earlier proposal along similar lines, Wyneken (1868; translated in Van Riper, 1982 p281) describes the will to speak during stuttering as “partially paralysed by doubt … and one which is directly opposed to the will proper”. Wyneken goes on to liken stuttering to “…when somebody, for example, wants to venture a jump, but in the very moment in which he leaps doubts that he will succeed. Often he can no longer stop the leap, but also does not jump with sufficient assurance, and so does not reach his goal.”

The approach-avoidance proposal is updated in several ways. Firstly, the core hypothesis of increased reflexivity for PWS corresponds to own speech having increased salience when interpreted through the auditory system. Secondly, it is proposed that the unconscious interpretation of own speech operates with the high degree of automaticity proposed for unconscious processes in dual process theory (e.g. as per Evans, 2007; Kahneman, 2011). The double approach-avoidance conflict proposed by Sheehan is thus fragmented between unconscious and conscious processes. The final proposal is that stuttering occurs at times when there is uncertainty about the message being delivered. The uncertainty might, for example, relate to message content (e.g. whether the message being conveyed is accurate) or to message appropriateness (e.g. whether the message should be delivered to a particular audience, or at a particular time). The uncertainty could also be learned (e.g. from previous experience with stuttering – this would account for the difficulty many who stutter have in saying their own name).

Putting all of these components together, the overall proposal is that whenever the speaker unconsciously interprets own speech with uncertainty, nerve signals are created which block the ongoing speech act. At the same time, the speaker notices difficulty and consciously generates nerve signals intended to continue the speech act. Articulatory muscles respond to both conscious and unconscious processes, and so simultaneously receive innervation which is consistent with completion and cessation of an utterance. The resultant activity is behaviourally observable as stuttering. Figure 7 summarises the activity diagrammatically.

**Figure 7:**
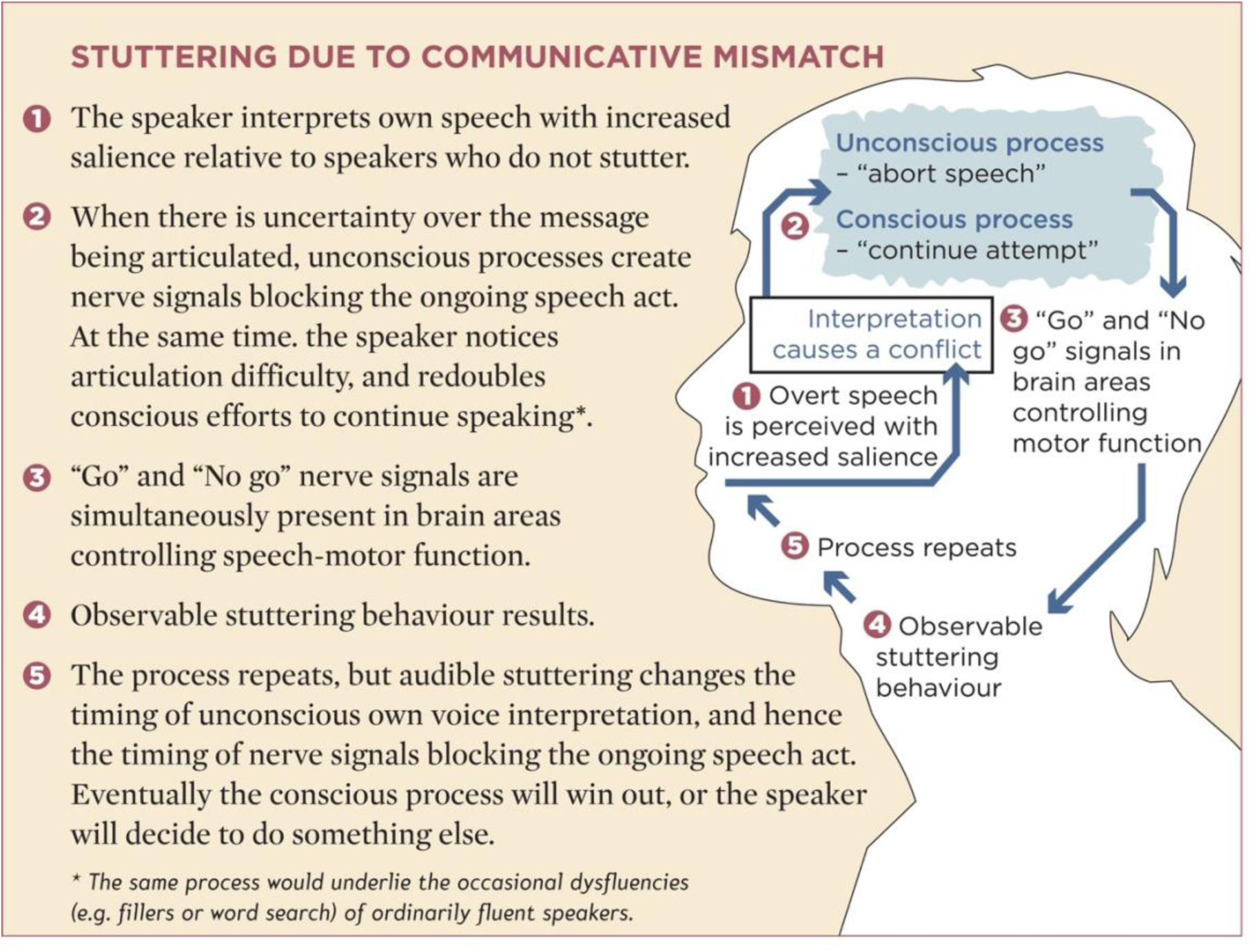
Stuttering due to communicative mismatch. This is based on the approach-avoidance conflict model of stuttering developed by Sheehan (1953, 1958, 1970, 1975), but updated to reflect contemporary understanding of unconscious processes. © Creative Commons 4.0 licence.

From a psycholinguistic perspective, REMATCH places stuttering at the semantic-pragmatic interface. In Levelt’s model (figure 6) stuttering would occur within the Conceptualiser. This differs from the psycholinguistic models in section 3.2.3.1, which place stuttering in the Formulator (or, for EXPLAN, between the the Formulator and Articulator). Section 3.4.1.2 will discuss a way to reconcile such psycholinguistic models within REMATCH.

### 3.4 Explanatory Power

#### 3.4.1 Biological considerations

##### 3.4.1.1 Neurological substrate

Systematic review of grey matter structural neuroimaging in adults who stutter (AWS) shows increased volume in the right superior temporal cortex and right precentral cortex compared to control groups of ordinarily fluent speakers (Etchell et al., 2017; see review for finer granularity and additional areas). These are homologues of areas in the left hemisphere thought to be important for speech and language (Hickok & Poeppel, 2007). Activation likelihood estimation (ALE) meta-analysis of diffusor tensor imaging shows AWS have reduced fractional anisotropy in the callosal body, and in dorsal white matter tracts connecting grey matter regions considered important for auditory and motor function (Neef et al., 2015; see review for finer granularity and additional areas). Possible interpretations of reduced FA include demyelination, larger axon diameter, lower packing density or increased axonal membrane permeability (Jones et al., 2013). Fractional anisotropy does not detail direction of information flow between grey matter areas.

ALE meta-analyses of functional neuroimaging in AWS during speech tasks show overactivation in areas corresponding to motor activity and underactivation in areas corresponding to auditory activity (Budde et al., 2014; Belyk et al., 2015; see state/trait comparisons in these meta-analyses for additional areas and finer granularity). Cerebellar vermis in AWS is underactive compared to controls, but overactive during stuttering. Neuroimaging of children who stutter shows differences from controls in several of the areas identified for adults who stutter (see, for example, Garnett et al., 2018; Kronfeld-Duenias et al., 2018; Koenraads et al. 2020), suggesting that the neurodevelopmental trajectory for stuttering diverges from that of ordinarily fluent speakers close to stuttering onset.

The core hypotheses of reflexivity and communicative mismatch will be traced through the brain areas just described. The route followed will describe a chronological sequence from audition of own voice, through cortical activity consistent with an ongoing speech act, to the creation of observable stuttering via speech-motor activity. References are as per the review articles already cited (Etchell et al., 2017, Neef et al., 2015, Belyk et al., 2015 and Budde et al., 2014) with further references introduced as necessary.

According to the Concurrency Hypothesis (Section 2), own voice will be identified through coincidence detection between cochlear and vestibular afferents. Only two studies have assessed the vestibular system in PWS. Langová et al. (1975) found that horizontal nystagmus evoked during speech is more pronounced in PWS than in controls using rotary chair testing. Gattie et al. (submitted) found the vestibular-evoked myogenic potential, an indirect functional test of the vestibular brainstem and periphery, is smaller in PWS than in controls. The suggestion is of divergence in central vestibular function, and/or the nature of conduction along the VIII cranial nerve, between PWS and controls.

Interpreted according to the Concurrency Hypothesis (section 2.3.2), a smaller vestibular input to the auditory brainstem would correspond to a lower likelihood for coincidence detection in cells whose excitation depends upon summation of synaptic input from multiple fibres (e.g. octopus cells in the cochlear nucleus, or principal cells of the medial superior olivary complex; Golding & Oertel, 2012). It follows that the ascending auditory stream at later stages of the ascending auditory pathway, or in temporal cortex, will be more weakly identified as an own voice stream in PWS than in controls. Inputs to cerebellar vermis will also be reduced during vocalisation for PWS (i.e. as per figure 3).

The sum of activity so far (smaller vestibular input to afferent streams of neural activity through the cerebellum and auditory brainstem) would more weakly identify own voice in PWS than in ordinarily fluent speakers. This occurs because the coincidence detection proposed by the Concurrency Hypothesis will be weaker with a smaller vestibular input. The weaker identification of own voice would in turn correspond to the increased reflexivity hypothesised for PWS. It is almost as if own voice is interpreted as for the voice of another speaker. From a systems control perspective (e.g. as per Jones et al., 2016), this would be referred to as inadequate sensory gating of the own voice auditory stream.

In the cerebrum, afferent own voice streams mediated via auditory brainstem and cerebellum could alter function in two brain areas which have repeatedly been identified as important in stuttering research. One of these is the cortico-basal ganglia-thalamo-cortical loop (Milardi et al., 2019), which is reviewed specifically in relation to stuttering by Chang & Guenther (this issue). The other area is temporal cortex, and in particular the temporo-parietal junction. Recall in this regard the discussion of section 2.4.1, that an own voice auditory stream would provide a target for the proposed efference copy of the speech plan in predictive feedback control models. Several authors have proposed that a difference in such moderation, or in auditory-motor mapping, between PWS and controls underlies the observed stuttering behaviour (Max et al., 2004; Brown et al., 2005; Hickok et al., 2011; Cai et al., 2012). The proposals have received little support in direct tests (e.g. Beal et al., 2010; Liotti et al., 2010). However, tests have used vocalisation versus AC playback protocols. As described in section 2.4.1, vocalisation versus AC playback protocols do not use a physiologically valid own voice stimulus in the playback condition, and as such do not evaluate speech-induced suppression. Accordingly, the proposal that moderation of temporal cortex by speech-motor activity differs between PWS and controls remains live. It is one of the possibilities for the hypotheses of concurrency and reflexivity when applied to stuttering via a predictive feedback control model. Investigation of the temporo-parietal junction is of particular interest, because it has repeatedly been identified as important for self-other distinction (Steinbels, 2016), and contains an area in the Sylvian fissure hypothesised as important for language control (Hickok, 2017). The role of the cerebellum may also be crucial. The cerebellum is repeatedly found to have involvement in speech perception (meta-analysis in Skipper & Lametti, 2021). This includes high level tasks involving semantics, grammar, and comprehension (Ackermann & Brendel, 2016; Mariën & Manto, 2015).

The hypothesis of communicative mismatch is based on approach-avoidance conflict (Sheehan 1953; 1958; 1970; 1975), which contemporary research (review in Aupperle & Paulus, 2010; Barker et al., 2019) places in the insula, amygdala, prefrontal cortex and the basal ganglia. All of these areas have been identified as showing a difference between PWS and controls in neuroimaging research (Yang et al., 2017; Toyomura et al., 2018; Budde et al, 2014; Etchell et al., 2017; see Garcia-Barrera & Davidow, 2015, for discussion of connection between prefrontal and anterior cingulate cortex in error monitoring). The basal ganglia in particular are crucial to the hypothesis of communicative mismatch. This is because conflict between selection and inhibition of competing actions, sometimes termed as “Go” and “No Go” (Mink, 1996; Bahuguna et al., 2015; Dunovan et al., 2015; Mink, 2018), could create involuntary muscular activity similar to that observed in stuttering. Thus, following the sequence described in this section from audition to articulation, basal ganglia activity would be the most proximal cause of observable stuttering behaviour (see Arenas, 2017, for a proposal emphasising functional importance of the subthalamic nucleus). Frontal and parietal cortex associated with speech-motor control would show state and trait differences in stuttering due their involvement in basal ganglia pathways (Albin et al., 1989; DeLong et al, 1990; Calabresi et al., 2014), and also due to white matter connection to temporal cortex important for auditory-motor integration (e.g. the efference copy proposed in speech-motor models), including commissural connection to homologues. The basal ganglia and cerebellum are interconnected via the thalamus (Hoshi et al., 2005; Bostan et al., 2010, Pelzer et al., 2017; Caligiore et al., 2017; Cacciola et al., 2017; Bostan & Strick, 2018) and are both proposed to have involvement in language processing and vocal learning (Booth et al., 2007; Pidoux et al., 2018). This underscores the prospective importance of the vestibular-cerebellar pathway (figure 3) for own voice identification in stuttering, and of cerebellar input to the cortico-basal ganglia-thalamo-cortical loop in stuttering (Chang & Guenther, this issue).

Dopamine levels in the basal ganglia affect action selection (Mink, 1996; Reynolds et al., 2001; Haber, 2014; Schultz, 2016), with differences in the dopamine system between PWS and controls found using positron electron tomography (Wu et al., 1997) and through pharmaceutical intervention (Maguire et al., this issue). The basal ganglia have been repeatedly identified as important in stuttering research (Alm, 2004; Metzger et al., 2017; Chang & Guenther, this issue).

##### 3.4.1.2 Stuttering subtypes

The neurological substrate described in the previous section encompasses almost the entire brain. This raises the possibility that preconditions for stuttering may require a difference from ordinarily fluent speakers in the function of not just one brain area, but several (ie. as per Ludlow & Loucks, 2003). Such a view underlies multifactorial models of stuttering (e.g. Smith & Kelly, 1997; Starkweather, 2002; Walden et al., 2012; Smith & Weber, 2017), which examine the interplay between genetic, organismic and environmental contributing factors.

One way to develop such models would be through subtyping stuttering. If there are discrete groupings of factors which contribute to stuttering, separation into such groupings prior to data analysis could enable more granular investigation and facilitate hypothesis formulation. Unfortunately, subtyping stuttering is difficult (see review in Yairi, 2007; Seery et al., 2007), largely due to the challenges of longitudinal data collection.

Subtyping is proposed here, based on the four track system of Van Riper (1973; 1982 p94–108). Symptomatology is identical to that of Van Riper, but the account is extended with the proposal that causation differs between tracks. Track I is proposed to correspond to stuttering developing as an isolated diagnosis. Track II corresponds to stuttering co-developing with at least one other diagnosis. Tracks III and IV are trauma-based, and may be psychogenic or neurogenic.

Track I would have a genetic basis and be based around increased reflexivity. The genetic basis may affect several brain areas (i.e. as per Ludlow & Loucks, 2003). For example, genetic investigation of stuttering has suggested that the nature of white matter may be integral to stuttering behaviour – lyosomal pathways or glial cells are implicated (Han et al., 2019; Benito-Aragón et al. 2020). PWS have reduced fractional anisotropy in dorsal white matter tracts which connect cortical regions having speech-motor and auditory function (Neef et al., 2015; Etchell et al., 2017). It would appear that genetic variations in stuttering might be connected to the structure of these dorsal white matter tracts. If so, it is not clear why the white matter structural variation should be focal to just these dorsal tracts (Watkins & Büchel, 2010; Drayna, 2010). One possibility is that genetic variation affects several white matter tracts. It may, for example, also manifest as reduced fractional anisotropy in the vestibular portion of the VIII cranial nerve. If so, the variation would be consistent with the finding of a weaker vestibular response in PWS by Gattie et al. (submitted), and would support interpretation according to the concurrency and reflexivity hypotheses presented in the current article.

This suggestion around genetics is just one example of a long-term investigative target for stuttering research. Many other possibilities pertain – not only variations within the neurological substrate described for stuttering in section 3.4.1.1, but moreover the interplay between genetic, organismic and environmental factors described in multifactorial models of stuttering. Investigation of which factors are necessary and/or sufficient for behavioural stuttering to manifest is a topic for ongoing research.

From this perspective, track II stuttering is a particular version of track I in which one of the variations contributing to the co-occurring diagnosis also contributes to stuttering behaviour. The existence of track II stuttering would explain why stuttering co-occurs with other diagnoses at a rate higher than chance. Depending on the nature of co-occurring diagnoses (and accepting that co-occurring diagnoses will sometimes remain pending) existence of track II stuttering could also explain why testing groups of PWS sometimes results in a subtle difference from controls in abilities such as executive function, language, reaction time and general motor control. The explanation would be that a co-occurring diagnosis, rather than stuttering, is causative of the test result. Such a distinction has been established in studies which split the PWS group into those with and without a co-occurring diagnosis (e.g. Cullinan & Springer, 1980; Liebetrau & Daly, 1981; McKnight & Cullinan, 1987; Kobayashi & Hayasaka, 2003)

REMATCH places stuttering at the semantic-pragmatic interface, but is compatible with the psycholinguistic models introduced in section 3.2.3.1. Application of the psycholinguistic models will help to distinguish tracks I–IV. Stuttering in tracks I and II could be described by a combination of EXPLAN (Howell, 2004; 2008) and the Variable Release Threshold hypothesis (Brocklehurst et al., 2013). It is suggested here that the account is updated such that it is REMATCH, rather than the compatible but less detailed anticipatory struggle hypothesis of Bloodstein (1975), which provides detail of the release threshold.

Tracks III and IV stuttering are proposed as having a trauma-based origin which could be psychogenic or neurogenic. When the trauma results in heightened self-awareness (Gallagher & Zahavi, 2021; Smith, 2020) increased attention to self-monitoring would follow. From the perspective of REMATCH the increased attention to self-monitoring would equate to increased reflexivity through a route other than the auditory system. Essentially, there is increased self-doubt about any speech act. In some cases the trauma could follow a profound emotional event (e.g. a bereavement or a family break-up), but it could also follow a more subtle series of events (Starkweather, 2002, lists possible environmental influencers on stuttering). Such stuttering could be described by the Vicious Circle Hypothesis (Vasić & Wijnen, 2005), in which monitoring of phonological error becomes hyper-vigilant. This type of stuttering could alternatively be explained from the perspective of EXPLAN and the Variable Release Threshold hypothesis. It would correspond to a release threshold which varies similarly to that of an ordinarily fluent speaker, but which is continuously subject to a multiplier greater than unity.

Other instances of tracks III and IV could be primarily caused by neural insult (e.g. transient ischemic attack, traumatic brain injury or neurodegenerative disease). If the effect of the neural insult is to alter function of a brain area important for phonological formulation, this type of stuttering could be described by the Covert Repair Hypothesis. However, neurogenic stuttering will be the most difficult to model. If the diagnosis is of a progressive neurological condition, stuttering may be transient prior to being masked by a wider range of symptoms involving language, speech-motor or executive function. In neurogenic stuttering with no other symptoms, behaviour may differ from tracks I–II due to the alteration in brain function having a random structural cause (neural insult) rather than proceeding through a genetic, developmental or psychological route.

Tracks I-IV may show overlap. For example, a child may have a genetic disposition to stuttering (track I) and experience environmental conditions creating psychosocial pressure (track III). This notion underlies the Demands and Capacities model (Adams, 1990; Starkweather & Gottwald, 1990; Starkweather, 2002), which is frequently interpreted as a genetic predisposition to stuttering becoming concrete following environmental influence. However, the predisposition need not be genetic; combinations of any of tracks I–IV, and/or single track etiologies, could just as well result in stuttering behaviour.

Developmental stuttering could involve any of tracks I–IV. Whereas absence of a plausible genetic or developmental contributory mechanism to stuttering in adulthood seems to limit adult onset stuttering to tracks III and IV. Thus, the rarity of adult onset stuttering (Ward, 2006, ch 16) is consistent with Van Riper’s (1973; 1982) finding that between 80–90% of his 300-strong caseload were tracks I or II. An exception would be adult onset where there is a history of childhood stuttering (Van Riper, 1982 p66). In such cases, reappearance may have psychogenic or neurogenic influence. For example, Shahed & Jankovic (2001) describe 12 persons who had stuttered in childhood but not as adults, and for whom stuttering reappeared following a diagnosis of Parkinson’s disease.

#### 3.4.2 Other explananda

The following sections address the explananda in tables 2 and 3, to which the reader might simultaneously refer.

##### 3.4.2.1 Linguistic and Situational

Variation in phonological formulation between stuttering subtypes was described in section 3.4.2. However, the main linguistic hypothesis within REMATCH is that own speech is interpreted in PWS with increased salience. Recall now the explananda in table 2. Unconscious processes are proposed to block an ongoing utterance whenever there is uncertainty about a speech act. Uncertainty is proposed to increase with propositionality, and hence stuttering will correlate with propositionality. Without an audience, a speech act cannot be performed. This explains why, unless PWS project an audience, stuttering will not occur when alone (Langová & Sváb, 1973). With authority listeners, even ordinarily fluent speakers experience increased salience when executing a speech act. For PWS, increased salience due to the authority listener combines with increased salience due to reflexivity, increasing the propensity for stuttering according to REMATCH.

These proposals are consistent with the observation of Sheehan (1958) that speech breakdown in stuttering coincides with the requirement “to say something important to someone important”. The proposals could be tested by following theoretical frameworks for pragmatics and social convention (e.g. Grice. 1957, 1989; Rescorla, 2019). For example, the exact loci of stuttered instances could be a project in experimental pragmatics (Noveck & Sperber, 2004; Meibauer & Steinbach, 2011; Noveck, 2018). Such a project might initially appear circular (stuttered phonemes are predefined as those with high propositionality). However, corpora of stuttered speech provide rich data, and can therefore be investigated following themes in pragmatics (e.g. Gricean implicatures, epistemic vigilance) using statistical techniques such as latent class or principal components analysis. Such an approach could also appraise changes in language use with development (e.g. within people who stutter there is a tendency for children to stutter on function words, and adults to stutter on content words). See also Eisenson & Horowitz (1945), Sheehan et al. (1967), Gould & Sheehan (1967) and MacKay (1969) for examples of work which could fit within a research programme for experimental pragmatics in stuttering.

##### 3.4.2.2 Anticipation, Consistency and Adjacency

Speakers can unconsciously scan ahead. This applies to spontaneous speech or when reading aloud. If message content scanned ahead is interpreted by the speaker according to REMATCH, the person who stutters will be able to predict when speech difficulty is imminent. In oral readings, uncertainty around any particular word is unchanged on repeated readings, because the underlying message has not changed. Therefore, stuttering has the same loci on repeat readings. When words are blotted out, the reader unconsciously anticipates what the word would have been (or infers intended meaning from the words remaining) leading to stuttering on the word that is unconsciously predicted to convey intended meaning to a listener. This will usually be an adjacent word to the word previously stuttered.

##### 3.4.2.3 Adaptation

Propositionality is reduced on repeat readings, since the listener is already aware of the message being delivered, and the speaker is aware of the message as well. Reduced propositionality in turn reduces salience. According to REMATCH, reduced salience will reduce the tendency for the speaker to unconsciously block an ongoing speech act. Essentially, reduced propositionality acts as a counter for the increased reflexivity proposed in PWS.

##### 3.4.2.4 Operant Conditioning

Speech-motor or psycholinguistic breakdown accounts of stuttering appeal to emotional or psychosocial stress to explain situational variation. A problem for such accounts is that they predict that stress should be very high in laboratory conditions with response-contingent stimulation (e.g. electric shock or time out upon stuttering), and therefore stuttering should increase. However, the converse is found: stuttering decreases with response-contingent stimulation.

The finding can be explained by REMATCH as response-contingent stimulation forcing an attentional shift in the speaker. The attentional shift is towards an increased conscious control of speech. This shift diminishes the influence of unconscious processing of speech, which according to REMATCH (figure 7) will reduce the amount of stuttering. Increasing conscious control is sufficiently effortful that unless an operant speaking technique such as fluency shaping has been learnt, PWS will not increase conscious control volitionally (see Constantino et al., 2020, for extended discussion). However, the continuous presence of response-contingent stimulation in laboratory conditions makes increased conscious control of speech unavoidable for the speaker.

##### 3.4.2.5 Alterations to audition during speech

Alterations to audition during speech will affect own voice identification according to the Concurrency Hypothesis (Section 2). Alterations which are effective are proposed to reduce reflexivity, and thereby to reduce stuttering according to REMATCH. The exact detail of audition changes effective for reducing stuttering is a topic for ongoing research. Timings in table 1 show a starting point. The effectiveness of long delays (e.g. 50 ms or more) may have more to do with phoneme or syllable recognition, or word recognition, than with own voice identification. If so, effectiveness of particular delay lengths will be variable, because the duration of word-initial phonemes is variable. The prediction from the Concurrency Hypothesis is that alterations on the time scale of a millisecond or less will be most effective. Such rapid alterations have not been tested other than by Howell et al. (1987), who showed that frequency shifts with a delay on the order of one millisecond were more effective at reducing stuttering than delays of 50 ms. Alterations most effective for reducing stuttering may depend on individual physiologies. If so, there is a prospect for tailoring the delay to individuals depending on EEG measurements. Such a project would be a part of, or be informed by, the investigation of own voice identification outlined in Section 2.

In alterations to audition involving a second speaker (shadowing or unison speaking) there is an additional benefit in that propositionality is also reduced (the second speaker is already aware of the message being delivered, and is encouraging delivery of that message). For this reason, unison speaking is the most effective way of reducing stuttering.

##### 3.4.2.6 Therapy effectiveness

REMATCH identifies the proximal cause of core stuttering behaviour as simultaneous “Go” and “No Go” signals in brain areas coordinating articulatory muscles, as described in section 3.4.1.1. Accessory and interiorised stuttering behaviours are explained in this regard as attempts by the speaker to resolve core stuttering behaviours (i.e. explanation as per Van Riper, 1982, ch 6–7).

Early stages of many stuttering therapies (e.g. the motivation, identification and desensitisation stages described by Van Riper, 1973) include psychological therapy, helping speakers to unlearn accessory and interiorised stuttering behaviours which have become engrained through habit. These early stages of stuttering therapy increase approach and decrease avoidance behaviours. For example, desensitisation therapy reduces emotionality attached to speaking situations. It is proposed that reduced emotionality will decrease the tendency to unconsciously block an ongoing speech act, and increase willingness to speak. The effect would be to reduce reflexivity, and thereby decrease stuttering according to REMATCH.

Speech work in therapies (e.g. the variation and adaptation stages in Van Riper, 1973) deliberately introduces prolongation to the beginning of syllables. Prolongation acts similarly to an alteration to audition during speech, and thereby reduces stuttering as described in section 3.4.2.5. An alternative strategy having the same effect would be to deliberately introduce repetition (Johnson. 1961). However, deliberate repetition is seldom used, perhaps because it is more noticeable than prolongation.

A major distinction between therapies is whether prolongation is on every syllable (fluency shaping) or only on syllables where stuttering is anticipated (block modification). See discussion in Ingham (1984, p328) or Gregory (1979). Prolongation on every syllable entails a continued attentional shift whilst talking. According to REMATCH, continued attentional shift will reduce stuttering (see section 3.4.2.4). Thus, fluency shaping has two methods reducing the amount of stuttering: syllable initial prolongation, and attentional shift. This would explain why fluency shaping programmes are often more effective than block modification programmes in reducing the amount of stuttering. However, fluency shaping programmes are effortful for the speaker (Constantino, 2020) and for this reason many PWS will prefer a block modification approach.

### 3.5 Discussion of the REMATCH hypothesis

The REMATCH hypothesis draws together breakdown and anticipatory struggle hypotheses of stuttering. In this sense, it is similar to, and compatible with, the Variable Release Threshold hypothesis of Brocklehurst et al. (2013). REMATCH goes into additional detail by specifying that the type of anticipatory struggle is an updated version of the approach-avoidance conflict proposed by Sheehan (1953; 1958; 1970; 1975). This update situates REMATCH in what Levelt (1989, 1999) refers to as the Conceptualiser. Thus, REMATCH is fundamentally different from (although compatible with) hypotheses which explain stuttering as breakdown in what Levelt refers to as the Formulator and/or Articulator. From this perspective, a major contribution of REMATCH is to provide a framework through which psycholinguistic and situational variation in stuttering can be investigated.

The updated approach-avoidance conflict in REMATCH is explained through a view of the unconscious proposed to be similar to that in dual process theory (e.g. as per Evans, 2007; Kahneman, 2011), and containing a high degree of automaticity. This provides a basis for investigation using cognitive science methodologies (e.g. as per the experimental pragmatics of Noveck & Sperber, 2004). The unconscious process in REMATCH could just as well have been explained as an update of repressed needs hypotheses of stuttering, in which the view of the unconscious is no longer necessarily that of psychoanalytic theory. This is possible because REMATCH contains a description of the moment of stuttering (figure 7) which can be compared to and informed by first person accounts. Thus, REMATCH promotes integration of qualitative and quantitive work in stuttering, and can furthermore provide a link to phenomenological accounts of stuttering (e.g. Ellis, 2020; Isaacs, 2020). Such integrations could inform psychological therapies for stuttering. They could also help to promote a social model of stuttering (Campbell et al., 2019), even within a world where neuroscientific research will remain within a medical model. Efforts in this direction are important if stuttering research is to be relevant to people who stutter.

## 4. General discussion

This article has described hypotheses of own voice identification and stuttering. The account has been highly detailed and with very broad scope because, as described in section 1, all of the hypotheses are proposed together as a best explanation argument. As such, it is necessary to show that the combined explanation has a high degree of explanatory power and parsimony.

The crux of this article is the Concurrency Hypothesis that own voice is identified through coincidence detection between the neural firing rates arising from deflection of cochlear and vestibular mechanoreceptors by the sound and vibration generated during vocalisation. Section 2 describes how the Concurrency Hypothesis provides a principled basis for self-environment distinction, with importance for considerations in cognitive science and philosophy of mind. The Concurrency Hypothesis was also applied to speech-motor research, in which it highlighted limitations in empirical support for the proposal that speech-motor activity modulates activity in temporal cortex. Finally, the Concurrency Hypothesis was applied to auditory scene analysis, in which it is proposed to provide the basis for a system of discrimination in multi-talker scenarios.

In Section 3, the Concurrency Hypothesis was developed into an explanation of stuttering. The initial step was to propose a quale, reflexivity. This refers to the phenomenology of hearing one’s own voice, and is proposed to differ between people who do and do not stutter. The account was then developed into an update of the approach-avoidance conflict model of stuttering (Sheehan 1953; 1958; 1970; 1975), referred to as REMATCH. This explains the moment of stuttering as a communicative mismatch. The speaker experiences own voice with increased salience, but this creates a mismatch whenever there is uncertainty about the ongoing message. In such cases, unconscious processes reinterpreting the message create nerve signals blocking the ongoing speech act, at the same time the speaker is consciously trying to continue. The resultant conflict is behaviourally observable as stuttering.

The Concurrency Hypothesis and REMATCH are core hypotheses. Many auxiliary hypotheses were introduced, mainly within the account of stuttering. These include the neurological substrate for stuttering, a proposal for subtyping stuttering, and a variety of process and contrastive explanations of data from stuttering research. These auxiliary hypotheses are likely to change with time, and are provided here as a snapshot so that the scope of the intended explanation of stuttering is apparent.

The Concurrency Hypothesis could be applied groups other than people who stutter, and who are expected to show differences from controls in own voice identification. Some examples of such groups include those experiencing auditory and/or vestibular neuropathy (Kaga, 2016) and those experiencing auditory hallucination (e.g. in schizophrenia – McLachlan, Phillips, Rossell & Wilson, 2013; Matthews et al., 2013; Weintraub et al., 2012; Waters & Fernyhough, 2019).

Testable predictions generated by the hypotheses in this article are described in sections 2 and 3. One of these predictions is that people who stutter should show a difference from controls in tests of the vestibular system. This was appraised by Gattie et al (submitted) with the finding that vestibular response is weaker in people who stutter than in paired controls. The result is consistent with the only prior research on the vestibular system in people who stutter (Langová et al., 1975) and supports the hypotheses presented in this article.

## 5. Conclusion

The major recommendation from this article is that researchers should use physiologically valid stimuli when investigating own voice in speech and language research. Using stimulation over air conduction only, even with a sound pressure level increase to perceptually match the loudness experienced during vocalisation, does not generate physiologically valid stimuli. Instead, stimuli should consist of a combination of air conducted sound and body conducted vibration which is binaurally symmetric, and has coincident arrival at both inner ears.

## Author Contributions

Conceptualization: ideas; formulation of the overarching research goals and aims

MG

Methodology: development or design of methodology or creation of models

MG

Software: programming; software development; designing computer programs; implementation of computer code or algorithms; testing code components

MG

Validation: verification of the replication and reproducibility of results, experiments, or other research outputs

MG

Formal analysis: application of statistical, mathematical, computational, or other techniques to analyse or synthesise data

MG

Investigation: conducting the research and investigation process, specifically performing the experiments or data collection

MG

Resources: provision of study materials, materials, instrumentation, computing resources, or analysis tools

MG

Data curation: annotation, scrubbing, or maintenance of research data (including software code, where it is necessary for interpreting the data itself)

MG

Writing—original draft: preparation, creation and/or presentation of the published work, specifically writing the initial draft

MG

Writing—review & editing: critical review, commentary, or revision

MG, KK, EL

Visualization: preparation, creation and/or presentation of the published work, specifically data presentation or visualisation

MG

Supervision: oversight and leadership responsibilities, including mentorship

KK, EL

Project administration: coordination of the research activity planning and execution

MG, KK, EL

Funding acquisition: acquisition of financial support

MG, KK

## Acknowledgments

Thanks to Paul Brocklehurst for comments on this article. Any shortcomings in the finished article are solely the responsibility of the named authors.

